# *S. cerevisiae* cells can grow without the Pds5 cohesin subunit

**DOI:** 10.1101/2022.05.21.492904

**Authors:** Karan Choudhary, Ziv Itzkovich, Elisa Alonso-Perez, Hend Bishara, Barbara Dunn, Gavin Sherlock, Martin Kupiec

## Abstract

During DNA replication, the newly created sister chromatids are held together until their separation at anaphase. The cohesin complex is in charge of creating and maintaining sister-chromatid cohesion (SCC) in all eukaryotes. In *S. cerevisiae* cells, cohesin is composed of two elongated proteins, Smc1 and Smc3, bridged by the kleisin Mcd1/Scc1. The latter also acts as a scaffold for three additional proteins, Scc3/Irr1, Wpl1/Rad61, and Pds5. Although the HEAT-repeat protein Pds5 is essential for cohesion, its precise function is still debated. Deletion of the *ELG1* gene, encoding a PCNA unloader, can partially suppress the temperature-sensitive *pds5-1* allele, but not a complete deletion of *PDS5*. We carried out a genetic screen for high copy number suppressors and another for spontaneously arising mutants, allowing the survival of a *pds5Δ elg1Δ* strain. Our results show that cells remain viable in the absence of Pds5 provided that there is both an elevation in the level of Mcd1 (which can be due to mutations in the *CLN2* gene, encoding a G1 cyclin), and an increase in the level of SUMO-modified PCNA on chromatin (caused by lack of PCNA unloading in *elg1Δ* mutants). The elevated SUMO-PCNA levels increase the recruitment of the Srs2 helicase, which evicts Rad51 molecules from the moving fork, creating ssDNA regions that serve as sites for increased cohesin loading and SCC establishment. Thus, our results delineate a double role for Pds5 in protecting the cohesin ring and interacting with the DNA replication machinery.

**IMPORTANCE:** Sister chromatid cohesion is vital for faithful chromosome segregation, chromosome folding into loops, and gene expression. A multisubunit protein complex known as cohesin holds the sister chromatids from S-phase until the anaphase stage. In this study, we explore the function of the essential cohesin subunit Pds5 in the regulation of sister chromatid cohesion. We performed two independent genetic screens to bypass the function of the Pds5 protein. We observe that Pds5 protein is a cohesin stabilizer, and elevating the levels of Mcd1 protein along with SUMO-PCNA accumulation on chromatin can compensate for the loss of the *PDS5* gene. In addition, Pds5 plays a role in coordinating the DNA replication and sister chromatid cohesion establishment. This work elucidates the function of cohesin subunit Pds5, the G1 cyclin Cln2, and replication factors PCNA, Elg1 and Srs2 in the proper regulation of sister chromatid cohesion.

## INTRODUCTION

Cohesin is a conserved protein complex that has two remarkable activities: i) it can tether two regions of chromatin (within the same DNA molecule or between DNA molecules) (1) and ii) it can extrude loops of chromatin (2, 3). These activities mediate sister chromatid cohesion (a mechanism that holds together the newly replicated DNA molecules from S-phase until anaphase) and facilitate condensation, DNA repair, and transcription regulation of a subset of genes (4). The temporal and spatial regulation of these cohesin-dependent biological processes is achieved in part by the complex regulation of cohesin. Identifying the modes of cohesin regulation and their coordination remains an important but elusive goal of the field.

In all eukaryotic organisms, including *S. cerevisiae*, the cohesin complex consists of four core subunits: two <u>S</u>tructural <u>M</u>aintenance of <u>C</u>hromosome (SMC) proteins, Smc1 and Smc3, one kleisin protein, Mcd1/Scc1 (hereafter referred as Mcd1) along with the HAWK family protein (HEAT proteins associated with kleisin) Scc3 [reviewed by (5)]. Various essential and non-essential proteins regulate cohesin life cycle. Here we focus on elucidating the function of Pds5, one of cohesin’s most critical and complex regulators. Pds5 is a HEAT repeat protein with no apparent catalytic activity that binds to Mcd1 near its N-terminus and plays central roles in cohesin function (6–8). Pds5 is important for human health as Pds5p deficiency has been linked to many cancers (9).

Pds5 was initially identified as a factor required for the maintenance of cohesion from S phase until the onset of anaphase (6, 10). The Pds5 protein is conserved and essential for cell division in almost all eukaryotes (4). However, subsequent studies have shown that Pds5 seems to regulate cohesion both negatively and positively. It is required for cohesion establishment and maintenance (6, 11). It also forms, with the Wpl1 protein, a complex that counteracts cohesion (12). How Pds5 plays such diverse and sometimes opposing roles in cohesin function? Several mechanistic studies have provided important clues.

SCC is a cell cycle-regulated phenomenon, and co-entrapment of sister DNA (establishment) is dependent on DNA replication. In *S. cerevisiae*, cohesin binding to chromatin starts in late G1; however, the cohesin rings are converted into cohesive structures only during DNA replication (13). The conserved acetyl-transferase Eco1 is essential for replication-dependent cohesion establishment (14, 15). Eco1 moves with the replication fork and acetylates the Smc3 protein at conserved lysine residues (K112, K113 in yeast) located in the head domain of Smc3 (16). Pds5 binding to cohesin enhances its acetylation by the Eco1 acetyl transferase (11). Also, Pds5 is known to block cohesin’s ATPase activity(17, 18) and antagonize the cohesin removal from the chromosomes by Wpl1 (11). However, other results contradict this Wpl1-centered view of the role of Smc3 acetylation and suggest that Pds5 binding to cohesin promotes SCC by a second, yet to be defined step (19).

In addition, Pds5 maintains cohesion, at least in part, by antagonizing the polySUMO-dependent degradation of cohesin (20, 21) and thereby stabilizing the complex. Pds5 binding to cohesin also promotes removal of unacetylated cohesin from chromosomes because Pds5 is a scaffold for Wpl1’s interaction with cohesin (12). However, many aspects of Pds5’s regulation of cohesin remain to be elucidated. The importance of Pds5 in blocking cohesin poly-SUMOylation was demonstrated by identifying mutations in SUMO and SUMO-modifying enzymes that suppress the inviability of Pds5 deficiency. However, other phenotypes of Pds5 deficiency were not suppressed (20–22) indicating that regulating the SUMO status of cohesin is only one function of Pds5.

PCNA, which recruits Eco1 to carry out its function, is a homotrimeric ring that plays a central role in DNA replication and repair. It acts as a processivity factor for the replicative DNA polymerases and as a “landing platform” on the moving replication fork. A conserved RFC-like complex that includes the Elg1 protein is in charge of PCNA unloading during Okazaki fragment processing and ligation [reviewed in (13, 23)]. Deletion of *ELG1* is not lethal but leads to increased recombination levels, as well as elevated levels of chromosome loss and gross chromosomal rearrangements (24). Human ELG1/ATAD5 plays an essential role in maintaining genome stability and acts as a tumor-suppressor gene (25). In the absence of the *ELG1* gene, PCNA accumulates on the chromatin, mainly in its SUMOylated form (26, 27). Mutants lacking Elg1 exhibit defects in SCC and are synthetic lethal with hypomorphic alleles of cohesin subunits (28). Thus, it is surprising that deletion of *ELG1* can suppress the temperature sensitivity (TS) of the *pds5-1* allele (29).

In this article we investigate the mechanisms by which cells can survive in the complete absence of Pds5. By carrying out genetic screens for suppressors of *pds5Δ elg1Δ* double mutants, we identify novel features Pds5 that inform on its integration with other cohesin regulators.

## RESULTS

### Screening for suppressors of the *pds5Δ elg1Δ* double mutant

Pds5 is essential for cohesion and cell viability in yeast (10, 30) and mammals (31). Thus, most studies in yeast take advantage of the *pds5-1* mutant, which can grow at the permissive temperature of 25°C, but does not grow at temperatures higher than 34°C (20, 29, 30). Previous studies revealed that a deletion of the *ELG1* PCNA unloader suppresses the temperature sensitivity of *pds5-1* mutant cells, allowing them to grow at higher temperatures (29). We confirmed this result (data not shown) and tried to test whether the lack of Elg1 could also suppress a total deletion of *PDS5*. We created a *pds5Δ elg1Δ* double mutant strain kept alive by the presence of a *URA3-marked* centromeric plasmid carrying the *PDS5* gene. This strain, however, was unable to form colonies on 5-FOA (5-Fluoroorotic acid) plates, which select for Ura-cells that have lost the covering plasmid (**Figure S1A**). We thus conclude that whereas the deletion of *ELG1* can suppress the *pds5-1* temperature-sensitive allele, which may still carry some residual Pds5 protein at high temperature, it cannot rescue the complete lack of Pds5 protein.

To better understand the interactions between Pds5 and Elg1, we performed two independent genetic screens looking for the suppressors of the *pds5Δ elg1Δ* double mutant. We looked for high-copy-number suppressors on the first screen, whereas in the second screen, we searched for spontaneous mutations in the genome that allowed the *pds5Δ elg1Δ* strain to survive without the covering plasmid.

### Pds5 ensures cell viability by enhancing the amount of Mcd1 in cohesin complexes

In our high copy number suppressor screen, we transformed a *pds5Δ elg1Δ* strain kept alive by the presence of a covering *URA3-PDS5-TRP1* plasmid with a yeast genomic library overexpressed from a 2-micron plasmid marked with a *LEU2* marker [the Yeast Genomic Tiling Collection (32) (**Figure 1A**)]. We searched for colonies able to grow in the absence of the covering plasmid. Since 5-FOA resistant colonies could also arise from mutations in the *URA3* gene carried on the plasmid, we identified Leu+, 5-FOA^r^ (Ura-), Trp-colonies, and isolated their library *LEU2*-marked plasmid (**Figure 1A**).

**Figure 1.**
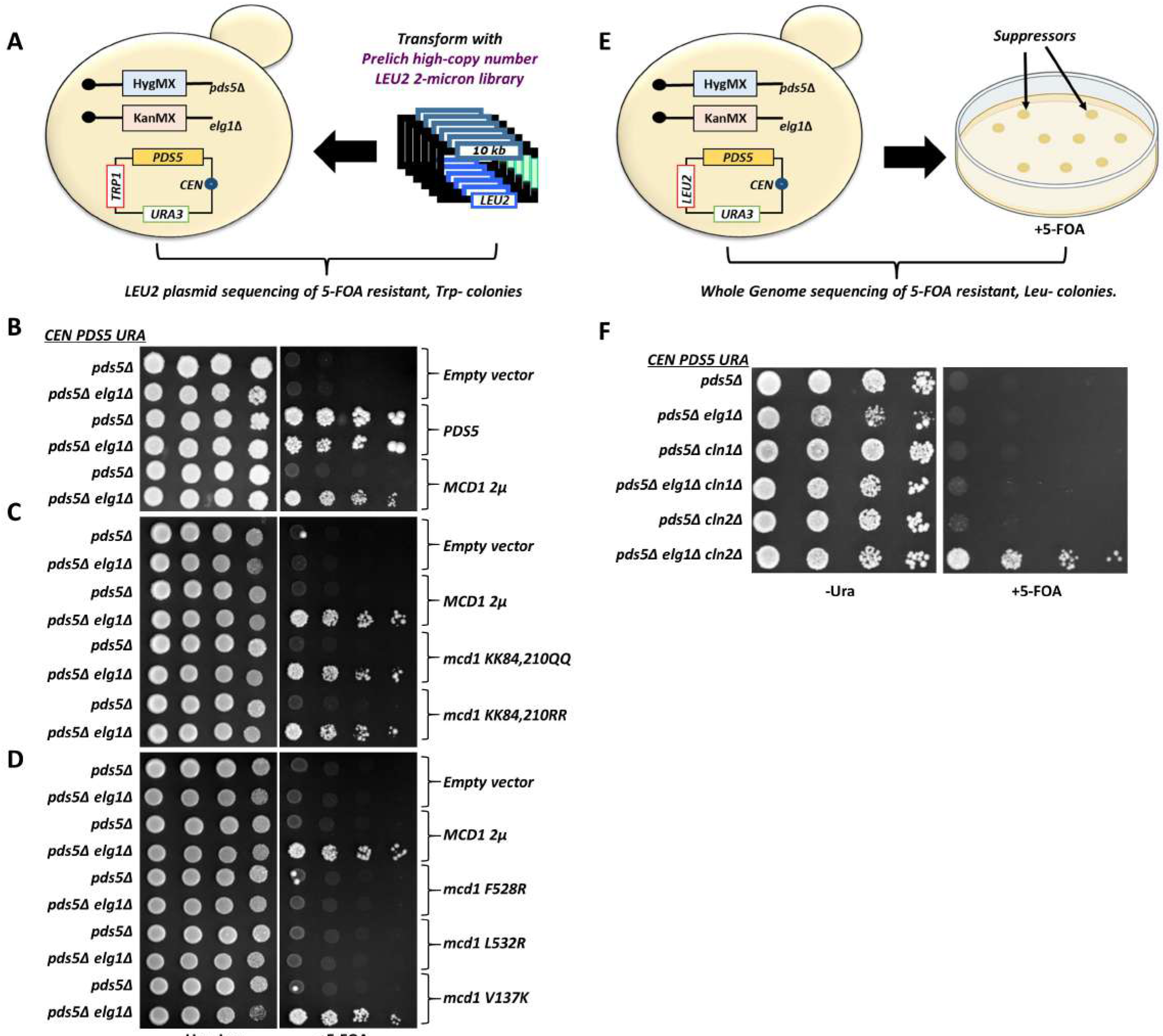
Screen for suppressors of the *pds5Δ elg1Δ* double mutant. **A)** Illustration of the experimental scheme for the high copy number suppressor screen. **B)** Fivefold serial dilutions of cells harboring either empty vector or high copy number vectors overexpressing *MCD1* or *PDS5* in addition to the covering plasmid (carrying the *URA3* and *PDS5* genes). **C), D)** Spot assay with fivefold serial dilutions of cells harboring either empty vector or high copy number plasmids overexpressing *MCD1* with different mutations at specified residues in addition to the covering plasmid (carrying the *URA3* and *PDS5* genes) **E)** Experimental regimen of a screen looking for the spontaneous suppressor mutants able to grow in the complete absence of *PDS5* and *ELG1*. **F)** Spot assay with fivefold serial dilutions spot assay of the *pds5Δ* background strains carrying specified gene deletions on –Ura and 5-FOA plates. All mentioned strains carry a Pds5 covering plasmid (carrying the *URA3* selection marker).

Out of the 80 Leu+ Ura-Trp-colonies obtained, 53 plasmids carried the genomic fragment carrying the *PDS5* gene, confirming the validity of our approach. Twenty-one additional plasmids carried a DNA fragment containing the *MCD1* gene. Mcd1 is one of the four core subunits of the cohesin complex. We further confirmed these results by transforming the cells with a subclone carrying only the *MCD1* gene. **Figure 1B** shows that overexpression of *MCD1* suppressed the lethality of *pds5Δ* in the absence of *ELG1*, but not in its presence.

To further understand the mechanism of this suppression, we selected different mutants of Mcd1 and observed their potential to rescue the lethality of *pds5Δ* and *pds5Δ elg1Δ* cells. We hypothesized that deletion of *ELG1* may elicit the DNA damage dependent, Chk1-dependent cohesion establishment pathway, which requires acetylation of Mcd1 at lysines 84 and 210 (33). If this proposition was true, then overexpression of the *mcd1-RR* allele (no acetylation possible) should not suppress, whereas overexpression of the *mcd1-QQ* (mimicking constant acetylation) should suppress the *pds5Δ elg1Δ* cells. However, both alleles were equally able to rescue the lethality of *pds5Δ elg1Δ,* suggesting that the rescue is independent of the DNA damage-mediated pathway **(Figure 1C)**. Furthermore, the deletion of the *CHK1* gene did not affect the suppression provided by Mcd1 overexpression (data not shown).

Overexpression of Mcd1 could be titrating an interacting protein; alternatively, it might be required to increase the levels of active cohesin. We thus introduced *MCD1* alleles unable to interact with cohesin (*mcd1-F528R* and *mcd1-L532R*) (34) or, as a control, an allele that does not interact with Pds5 (*mcd1-V137K*)(35). **Figure 1D** shows that only overexpression of the *mcd1* alleles that could be incorporated into the cohesin complex allowed the *pds5Δ elg1Δ* double mutant to grow on 5-FOA plates, ruling out a titration effect. The overproduction of different Mcd1 alleles was also confirmed by western blot in *pds5Δ* and *pds5Δ elg1Δ* double mutant background (**Figure S1B)**. Thus, increased levels of Mcd1 at chromatin allow *pds5Δ elg1Δ* to grow. The fact that overexpression of Mcd1 cannot suppress the single *pds5Δ* mutant but efficiently suppresses the double *pds5Δ elg1Δ* suggests that in the absence of Pds5, two independent changes are necessary: on the one hand an elevation of Mcd1 levels, on the other hand, something that the absence of *ELG1* is providing. Each of these two changes is by itself insufficient to allow *pds5Δ* strains to grow.

### Spontaneous mutations in the G1 cyclin *CLN2* ensure cell viability of *pds5Δ elg1Δ* double mutant

In our second screen, we looked for spontaneous mutants that allow the *pds5Δ elg1Δ* double mutant strain to lose its covering plasmid. We plated a large number of yeast cells on 5-FOA plates in several batches and looked for colonies that grew on 5-FOA plates and were Leu-. We confirmed that these colonies had lost the covering plasmid and performed whole genome sequencing to identify the suppressor mutations in the genome (**Figure 1E**).

Out of the 40 independent 5-FOA resistant, Leu-mutants that lost their covering plasmid, 23 carried *de novo* mutations in the *CLN2* gene. Most of the mutations were nonsense, frameshift, or indel mutations that inactivated the gene (**Figure S1C**). The *CLN2* gene encodes a G1 cyclin that is necessary for the transition between G1 and S phases. In order to test these results, we made a genomic deletion of *CLN2* gene in the *pds5Δ elg1Δ* background. As expected, the strain carrying the triple deletion of *pds5Δ elg1Δ cln2Δ* grew well on 5-FOA plates, suggesting that the *CLN2* deletion suppresses the lethality of the *pds5Δ elg1Δ* strain (**Figure 1F**). A second G1/S cyclin gene, *CLN1,* has 57% sequence identity (72% in the N-terminal region) to *CLN2* gene (36) and is expressed with similar timing, attaining maximal expression during the G1/S transition (37). Therefore, both *CLN1* and *CLN2* genes are considered functionally redundant (38). **Figure 1F**, however, shows that a deletion of *CLN1* could not suppress the lethality of the *pds5Δ elg1Δ* double mutant strain. As in the case of *MCD1* overexpression, the deletion of *CLN2* only allows growth of the *pds5Δ* strain if *ELG1* is deleted too, confirming the existence of two different pathways that need to be modified to allow life in the absence of Pds5.

### Pds5 counteracts mechanisms that limit Mcd1 levels in cells

Based on the results from our genetic screens, our working hypothesis was that the deletion of *CLN2* mimics the overexpression of *MCD1,* increasing its protein level. In the following experiments, we used an auxin-inducible degron (AID) in order to be able to degrade Pds5 conditionally. The AID-*PDS5* strain grew normally and showed no cohesion or cell cycle defects. Adding auxin to the medium leads to the rapid degradation of Pds5 (**Figure S2A, B).** We arrested the cells in the cell cycle at the M phase with nocodazole and treated them with auxin for 2 hours. As expected from previous studies (20), there is a significant decrease in the level of Mcd1 protein in the AID-*PDS5* strain compared with the untagged strain in the presence of Auxin (WT vs. *AID-PDS5,* p value=0.02) (**Figure 2A and B, S2C**). AID-*PDS5 elg1Δ* and AID-*PDS5 cln2Δ* strains treated with auxin showed a similar decrease of Mcd1 protein (WT vs. AID-*PDS5 elg1Δ* p value=0.01; WT vs. AID-*PDS5 cln2Δ* p value=0.02). Mcd1 levels, however, were improved in the AID-*PDS5 elg1Δcln2Δ* strain in the presence of Auxin (AID-*PDS5* vs. AID-*PDS5 elg1Δcln2Δ* p value=0.005) (**Figure 2A and B**). To follow the kinetics of Mcd1 protein in the absence of Pds5, we induced the degradation of Pds5 by adding auxin to mid-log cultures and then measured the level of Mcd1 every 20 minutes. Following Pds5 degradation, the Mcd1 protein levels significantly drop in the AID-PDS5 strain and in the single *elg1Δ* and *cln2Δ* mutants. In contrast, we observed a much slower kinetic of Mcd1 reduction in the AID-*PDS5 elg1Δ cln2Δ* mutant, which retained more than half of the Mcd1 protein levels after two hours of auxin addition **(Figure 2C-F)**. We conclude that only the concomitant deletion of *ELG1* and *CLN2* can restore enough Mcd1 to allow cell growth without Pds5.

**Figure 2.**
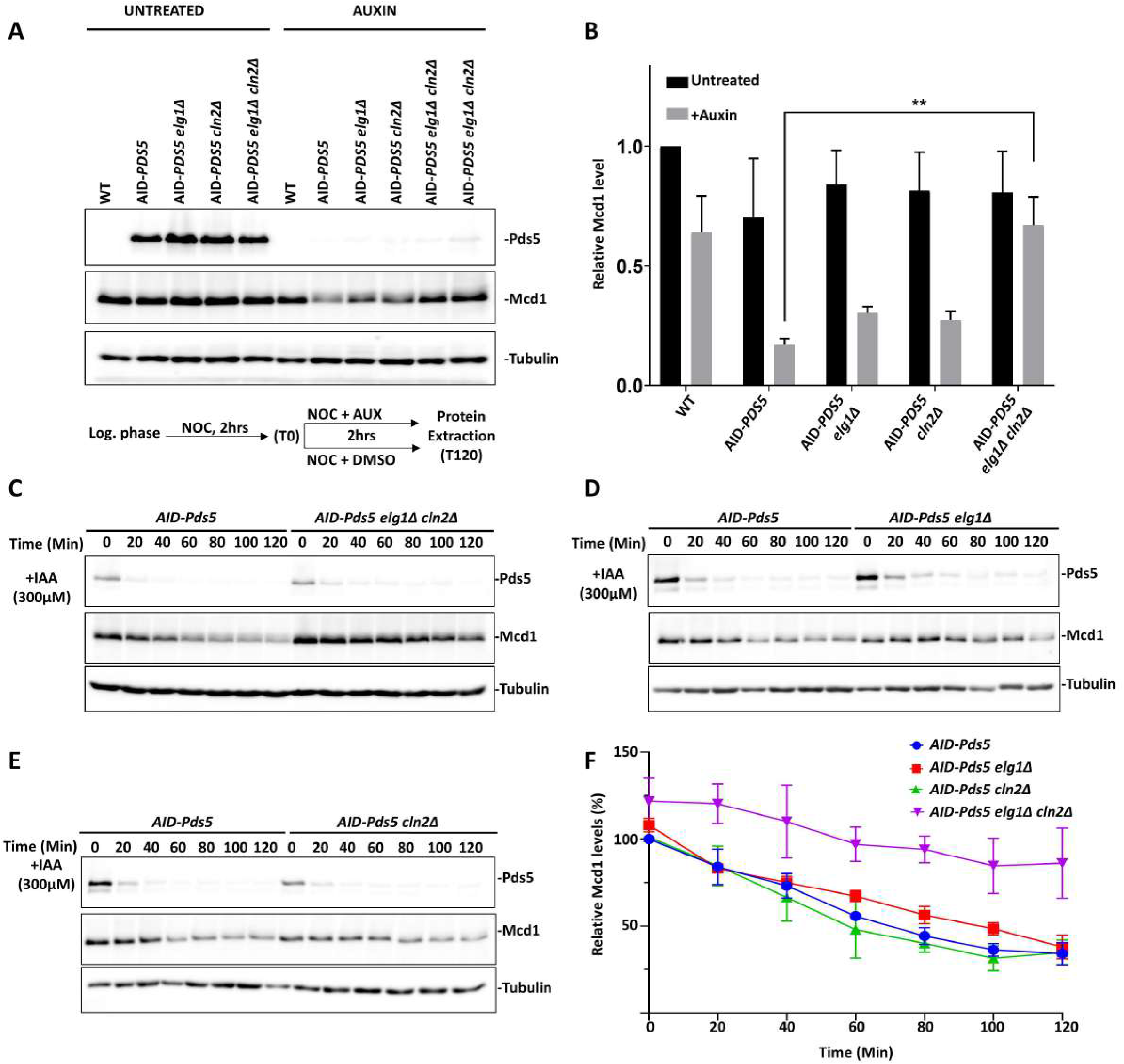
Deletion of *ELG1* and *CLN2* restores the Mcd1 protein level in the absence of Pds5. **A)** Western blot showing the Mcd1 protein level in different *PDS-AID* strains. Cells were harvested after arresting them in the G2/M phase by treatment with nocodazole (15μg/ml) for 2h, followed by the treatment with auxin (IAA, 300 μM). The experimental scheme is represented below the western blot panel. Mcd1 was probed with an anti-Mcd1 antibody, Pds5 was detected using anti-V5, and Tubulin was used as a loading control. **B)** Mcd1 protein levels normalized to those of Tubulin (mean ± SD; n=3). T-test analysis p-value ** ≤ 0.01. **C-E)** Western blot for the auxin-chase experiment. The cells of the indicated strains were grown until the log phase (time 0) and then treated with Auxin (IAA, 300 μM). Samples were taken every 20 minutes until completing a 2 hours experiment. **F)** Relative levels of Mcd1 protein normalized to those of Tubulin used as a loading control (n=3; % mean ±SD).

### *CLN2* deletion leads to overexpression of the Mcd1 gene

The high level of Mcd1 could be due to increased gene expression or to protein stabilization. To test whether the deletion of both *ELG1* and *CLN2* prevented Mcd1 degradation, we measured the half-life of Mcd1 in the presence of cycloheximide (CHX), which inhibits global protein synthesis. No significant difference in the rate of degradation was found between AID-*PDS5* and AID-*PDS5 elg1Δ cln2Δ* strains in the presence or absence of auxin (**Figure S3A-F**). Therefore, the increased levels of Mcd1 in the AID-PDS5 *elg1Δ cln2Δ* strain are not due to the increased stability of the Mcd1 protein. We thus hypothesized that the higher Mcd1 levels would be a consequence of increased Mcd1 transcription.

To test this hypothesis, we constructed a plasmid vector carrying short-lived GFP under the control of the *MCD1* promoter and a mCherry gene under the control of a constitutive *ADH1* promoter, which serves as an internal plasmid copy number control **(Figure 3A)**. We introduced this plasmid into the different AID-PDS5 strains and, using a flow cytometer, we measured the mean fluorescence intensity (MFI) for GFP and mCherry. We observe that the GFP/mCherry MFI ratio is significantly higher in AID-PDS5 *elg1Δ cln2Δ* and AID-PDS5 *cln2Δ* strains compared to AID-PDS5 in the absence or presence of Auxin **(Figure 3B)**. To validate the results from flow cytometry, we did a western blot analysis to observe the GFP protein levels in different strains carrying the reporter plasmid. In agreement with the earlier experiment, we observe a significant increase in the GFP protein levels in the AID-PDS5 *elg1Δ cln2Δ* and AID-PDS5 *cln2Δ* strains **(Figure 3C, D)**

**Figure 3.**
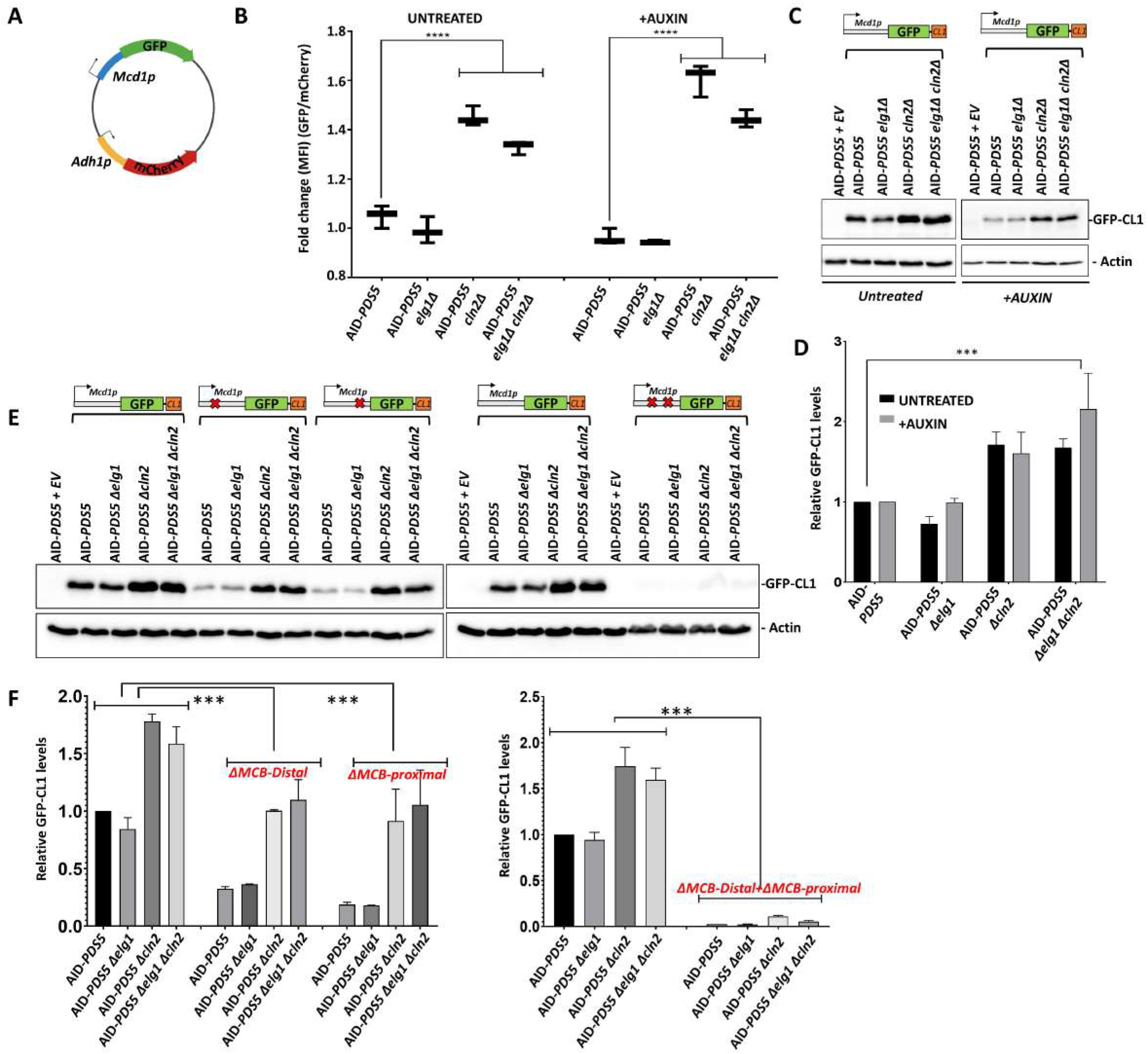
Mcd1 is overexpressed in *elg1*Δ cln2*Δ* double mutants. **A)** GFP-RFP plasmid with a short-lived GFP gene under the control of the Mcd1 promoter and internal control mCherry under the control of ADH1 promoter. **B)** Mean fluorescent intensity GFP/mCherry ratio from flow cytometery for different strains treated with Auxin (IAA 300μM for 2hrs (Right) and without Auxin (Left) (20,000 events, n=3). One way Anova p value *** ≤ 0.001. **C)** Western blot (anti-GFP) monitoring the GFP fused to CL1 degron protein levels in different strains expressed from a 2μ plasmid. Actin was used as a loading control. **D)** Western blot quantification of GFP levels normalized to the loading control actin. (Mean ± SD; n=3). T test analysis p value *** ≤ 0.001 **E)** Western blot (anti-GFP) monitoring the GFP-CL1 fusion protein levels expressed from a construct carrying a mentioned deletion in the MCB box in Mcd1 promoter. **F)** Western blot quantification of GFP levels normalized to the loading control actin. (Mean ± SD; n=3). One way Anova p value *** ≤ 0.001

Next, we wanted to understand how deletion of *CLN2* results in hyper-transcription of the *MCD1* gene. Cln2 is a G1 cyclin that promotes MBF-dependent transcription of many DNA replication and repair-associated genes during the G1-S phase transition (39). These genes contain distinct DNA binding domains for the MBF complex in their promoter (MCB motifs). The *MCD1* promoter contains two putative MCB motifs. Simultaneous deletion of both MCB motifs from the *MCD1* promoter completely abolished the GFP expression of all strains **(Figure 3E, F)**. These results show that the increased transcription of *MCD1* observed in *cln2Δ cells* is dependent on the MBF complex. Thus, the deletion of *CLN2* hyper-activates the MBF complex. Our results are consistent with previous studies, which also observed a high transcription of the MBF regulon in *cln1Δ cln2Δ* strain background (40, 41).

### Simultaneous deletion of *CLN2* and *ELG1* restores SCC to cells lacking Pds5

In the absence of Pds5, yeast cells die due to SCC defects. These cells are defective both in the establishment and maintenance of cohesion (30, 42). Similarly, *elg1Δ* strains were shown to be slightly defective SCC and exhibit increased levels of premature sister chromatid separation (28), although it was unclear whether the defect resides in the establishment or the maintenance of the cohesion. The simultaneous deletion of *ELG1* and *CLN2* provides robust growth in the absence of Pds5. To test whether SCC was also restored, we used the two-dot GFP assay (43). In this assay, an array of Lac operators is inserted in the chromosomal arms, recognized by a Lac repressor-GFP fusion protein. The binding of LacI-GFP to chromosomal arms can be observed under the fluorescent microscope as a bright dot in living yeast cells. When sister chromatids are adequately aligned by cohesion, only a single dot is seen, whereas two dots are observed in cells exhibiting premature separation (43).

We carried out a cohesion assay by synchronizing the cells in G1 with alpha-factor, then releasing the cells into the cell cycle in the presence of auxin and nocodazole (**Figure 4A, B**). This assay mainly measures the cells’ ability to establish functional cohesin molecules at the beginning of the S-phase. Under these conditions, the *AID-PDS5* strain exhibited more than 40% of cells with double dots, consistent with previous reports (20, 42). Deletion of *ELG1* or *CLN2* reduced the number of cells with premature sister chromatid separation, and the number was significantly further reduced in the AID-*PDS5 elg1Δ cln2Δ* strain (p value=0.021), indicative of an additive effect of the *elg1Δ* and *cln2Δ* mutations. As expected, no precocious chromatid separation was detected when auxin was omitted from the assay.

**Figure 4.**
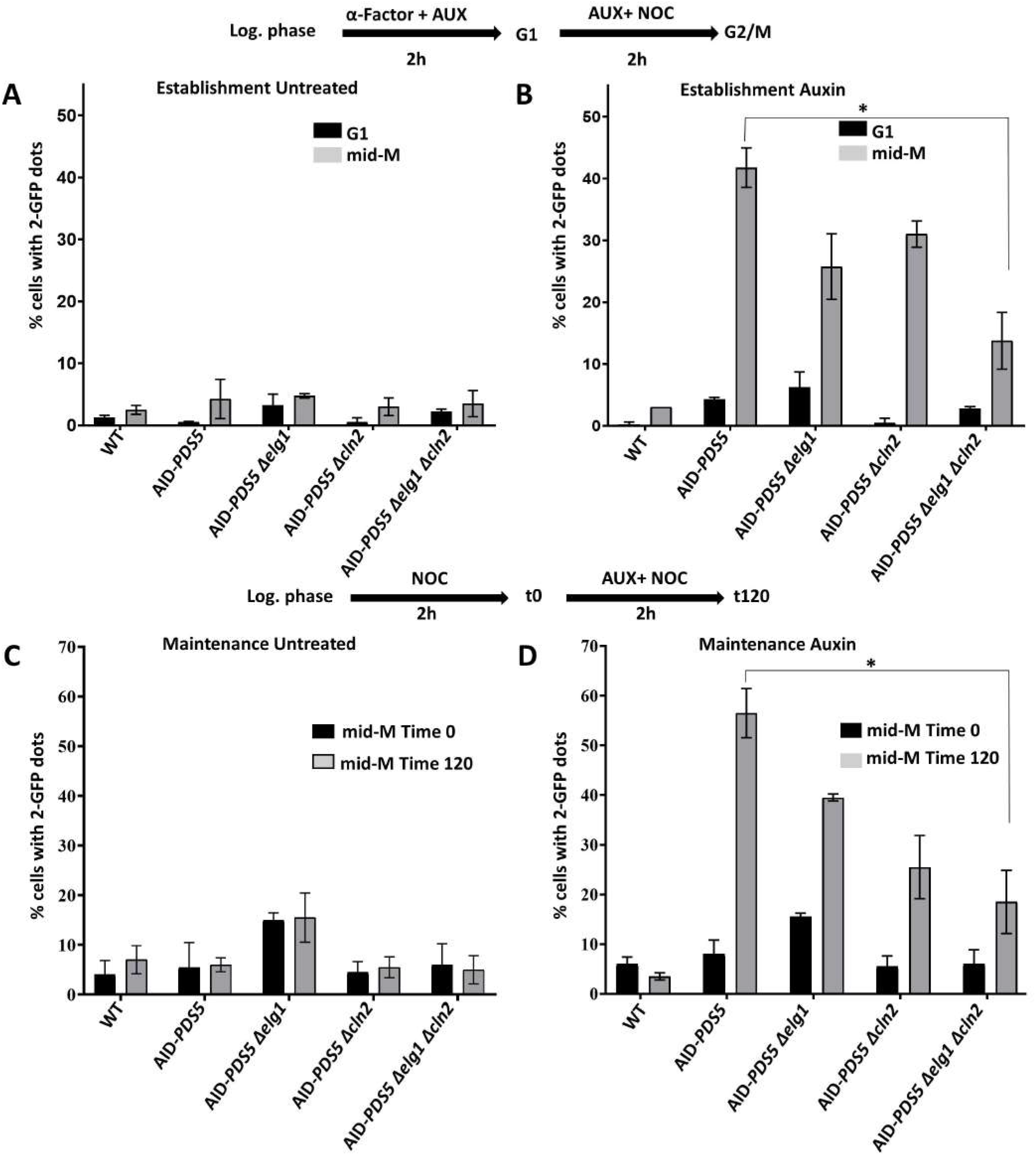
Deletion of *ELG1* and *CLN2* restores the sister chromatid cohesion defects in the absence of Pds5. **Cohesion establishment analysis**: Top panel-Experimental scheme for the cohesion establishment assay. **A)** Percentage of cells with 2 dots in mid-M phase without auxin treatment (n=3 with >200 cells per strain and experiment; mean ± SD) **B**) Establishment assay for auxin treated cells. α-Factor: **50** ng/mL; NOC: nocodazole 15μg/ml; PRON: pronase E 0.1 mg/ml. **Cohesion maintenance analysis** Mid panel-Experimental scheme for cohesion maintenance assay with AUXIN (IAA, 300 μM). The untreated experimental process was the same but without auxin. **C)** Percentage of cells with 2 dots for every strain without auxin treatment (n=3 with 200 cells per strain and experiment; mean ± SD) **D)** maintenance assay for auxin-treated cells in different strains. NOC: nocodazole 15μg/ml.

SCC is established during DNA replication in S-phase and maintained until anaphase. To test for SCC maintenance, cells were synchronized in early mitosis with nocodazole (after establishing cohesion) and maintained for 2 hours in the presence of auxin and nocodazole (**Figure 4C, D**). The AID-PDS5 strain exhibited a substantial maintenance defect: close to 60% of the cells exhibited two dots, consistent with previous reports (22). In this assay, the deletion of *ELG1* had only a minor effect, reducing the number of two-dot cells to ~40%. In contrast, the AID-*PDS5 cln2Δ* strain strongly reduced the number of cells with two dots, not significantly changed in the AID-*PDS5 elg1Δ cln2Δ* strain (T-test p-value = 0.022).

Our results thus point at two different roles of the *CLN2* and *ELG1* in sister chromatid cohesion: whereas both of them affect the establishment by separate pathways (and thus the mutants show additivity), the *elg1Δ* mutant plays only a small role once the sister chromatid cohesion has been established, whereas *cln2Δ* affects maintenance too. Both mutations are required for full viability (**Figure 1**).

### Elg1 contributes to the suppression by accumulating more PCNA on chromatin

The absence of Elg1 causes an accumulation of PCNA on the chromatin (44, 45). This increased level of PCNA is held responsible for most genome instability phenotypes exhibited by *elg1Δ* (46). To understand the function of Elg1 in SCC, we compared *pds5Δ cln2Δ* strains carrying a *URA3-PDS5*-covering plasmids, bearing different *ELG1* alleles in their genomes. The ability of the different alleles to provide Elg1 function was assayed by plating on 5-FOA plates **(Figure 5).** Whereas cells carrying an empty vector can lose their covering plasmid and grow on 5-FOA plates, the presence of the WT *ELG1* gene prevents growth, confirming our previous observations (**Figure 5A**). We observe that mutations in the *ELG1* Walker A motif, alleles with reduced ability to unload and recycle PCNA, such as *elg1-TT386,7DD, elg1-sim+TT386,387DD* (46), the Walker B mutant *elg1-DVD to KVK,* and the Walker A/Walker B double mutants (47) were unable to complement the *ELG1* deletion, and grew on 5-FOA plates. In contrast, mutations that do not greatly affect PCNA unloading, such as the *elg1-KK343,344AA* allele, fully complemented the Elg1 defect and thus were unable to allow growth on 5-FOA plates. A good correlation was observed between the degree of sensitivity to methyl methanesulfonate (MMS) [which reflects the amount of PCNA on the chromatin (46)] and the ability to lose the covering plasmid (**Figure 5A**). Moreover, PCNA variants that spontaneously disassemble from the chromatin (such as *pol30-D150E, E143K* or *S152P* (48), suppress the sensitivity of *pds5Δ elg1Δ cln2Δ* strains to MMS and prevent growth on 5-FOA (**Figure 5B**), indicating that the effect conferred by the deletion of *ELG1* is due to the increased levels of PCNA on chromatin. PCNA acts as a binding platform for the cohesin acetyltransferase Eco1 (16). Therefore, a simple hypothesis to explain the increased SCC in *elg1Δ* strains is that high levels of PCNA accumulation on chromatin caused by the *ELG1* deletion might elevate the chromatin levels of Eco1 protein. To test this possibility, we monitored Eco1’s overall chromatin abundance. We observe that although *elg1Δ* has higher levels of PCNA on chromatin, a corresponding increase in Eco1 abundance is not observed (**Figure 5C, D**).

**Figure 5.**
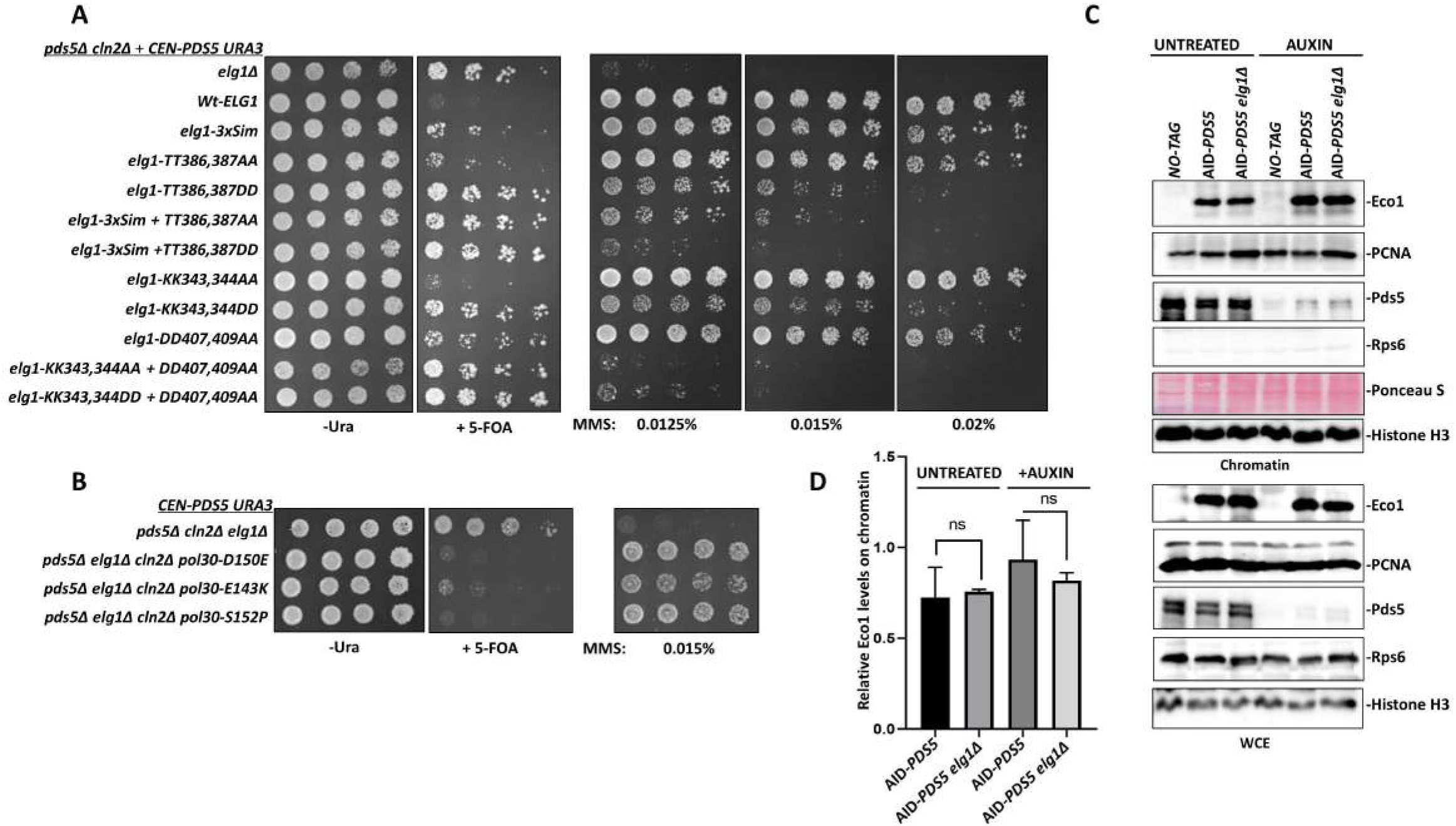
PCNA accumulation on chromatin promotes sister chromatid cohesion in the absence of Pds5: **A)** Spot assay with fivefold serial dilution of *pds5*Δ *cln2*Δ +(*CEN PDS5 URA*) strain carrying different mutants of Elg1 at the *ELG1* locus in the genome; on 5-FOA medium and plates containing the DNA damaging agent MMS at mentioned concentration. **B)** Spot assay with fivefold serial dilution of *pds5*Δ *cln2*Δ *elg1*Δ +(*CEN PDS5 URA*) background strain harboring disassembly prone PCNA mutations in the genomic copy of the *POL30* gene on 5-FOA plates. **C)** Chromatin Fractionation experiment showing the Eco1-3HA levels on chromatin in untreated and auxin (2hrs) - treated samples. Histone H3 was used as a chromatin marker and loading control, Rps6 was used as a cytoplasmic marker. **D)** The Graph represents the western blot quantification of the relative abundance of Eco1 protein on chromatin. (Mean ± SD; n=3). Student’s t-test ns= non-significant.

### Suppression of Pds5 depletion suggests that cohesin function is limited by Elg1-dependent removal of SUMOylated PCNA from DNA

The post-translational modifications of PCNA play an essential role in genome stability by coordinating several replication-coupled DNA damage tolerance pathways. When a replisome encounters a DNA lesion on a template strand, it may undergo modifications to activate a specific DNA damage bypass pathway [reviewed in (23)]. The Rad6/Rad18 dependent PCNA mono-ubiquitination at the K164 residue results in recruitment of an error-prone TLS (translesion synthesis polymerase) which adds more or less random bases at the damage site, allowing its bypass. The Rad5-dependent poly-ubiquitination at the K164 residue promotes an error-free template switch pathway (49). Similarly, PCNA SUMOylation at K127 and K164 by the SUMO ligase Siz1 recruits the helicase Srs2, which acts as a local anti-recombination factor (50).

In order to test whether PCNA modification plays any role in the suppression via *elg1Δ,* we mutated the conserved lysine residues K164 and K127 to the unmodifiable residue arginine in the background of *pds5Δ elg1Δ cln2Δ.* Interestingly, we find that PCNA mutations *pol30-K164R* or *pol30-KK127,164RR* both prevent plasmid loss and render cells inviable on FOA plates (**Figure 6A**).

**Figure 6.**
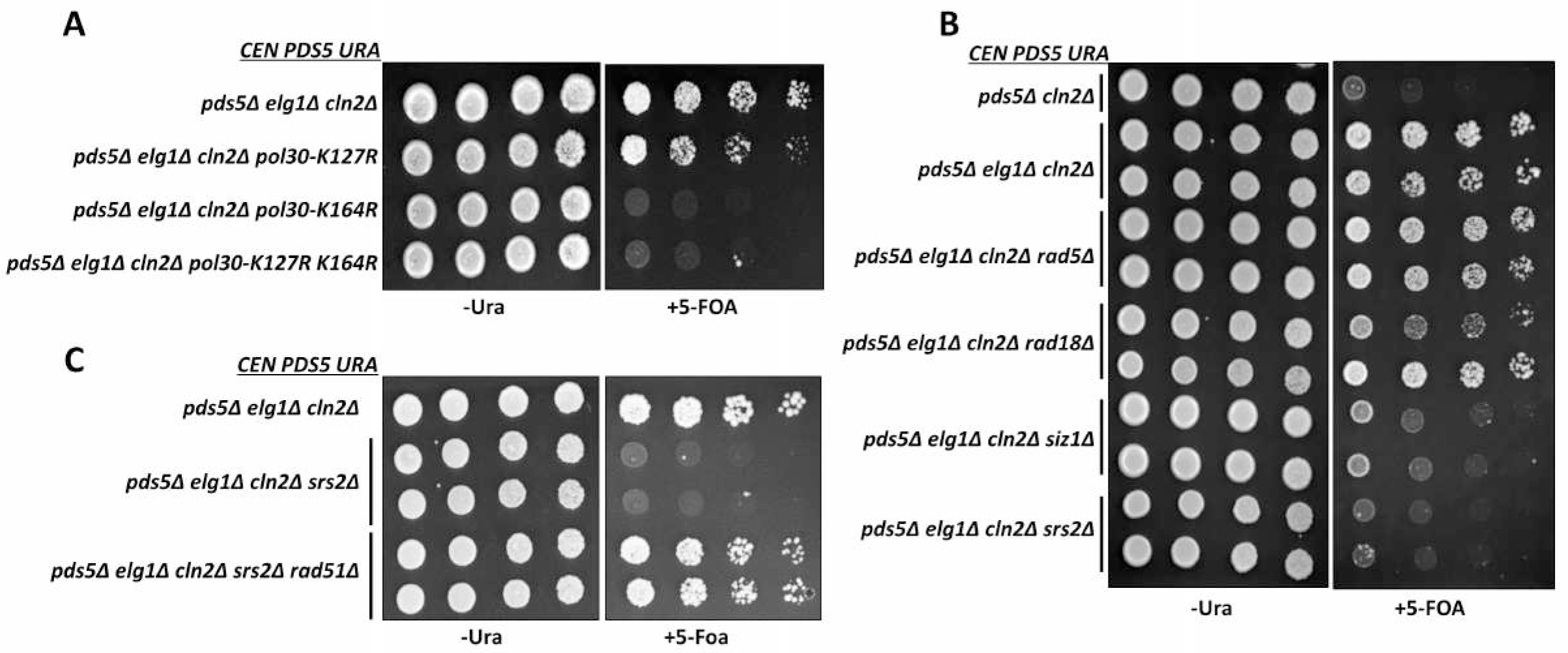
Sumo-PCNA accumulation on chromatin and Srs2 promote sister chromatid cohesion in absence of Pds5: **A)** Spot assay with fivefold serial dilution of *Pds5*Δ *cln2*Δ *elg1*Δ +(*CEN PDS5 URA*) background strain harboring point mutations at the key Lysine residue in the genomic copy of the *POL30* gene, on 5-FOA plates. **B)** Spot assay with fivefold serial dilution of *pds5*Δ *cln2*Δ *elg1*Δ +(*CEN PDS5 URA*) background carrying deletion of genes involved in PCNA ubiquitination (Rad5, Rad18) or PCNA SUMOylation pathways (Siz1) and the SUMO-PCNA interactor Srs2, on 5-FOA plates. **C)** Fivefold serial dilution of *pds5*Δ *cln2*Δ *elg1*Δ *srs2*Δ *rad51*Δ + (*CEN PDS5 URA*) and control strains on 5-FOA plates.

These results suggest that *elg1Δ* contributes to suppression by accumulating modified PCNA on chromatin. Next, we asked which kind of PCNA modification (SUMOylation or ubiquitination) is essential for promoting cohesion via *elg1Δ.* Deleting *RAD18* or *RAD5* in the *pds5Δ elg1Δ cln2Δ* background renders these strains susceptible to DNA damaging agent MMS; however, the lack of these factors did not affect the growth of yeast cells on FOA plates. In contrast, the deletion of the SUMO ligase Siz1 in the *pds5Δ elg1Δ cln2Δ* background abolished the rescue, and cells could not grow on FOA plates (**Figure 6B**). Therefore, we conclude that *elg1Δ* promotes cohesion by accumulating SUMOylated PCNA on the chromatin.

### Suppression of Pds5 depletion suggests that cohesin function is limited by Srs2-dependent removal of Rad51

Srs2 is an helicase that inhibits homologous recombination by stripping Rad51 filaments from the ssDNA (51). Srs2 binds to SUMOylated PCNA, and we have shown that *elg1Δ* strains accumulate a high level of both SUMOylated PCNA and Srs2 on chromatin (45). Based on this information, we deleted *SRS2* in *the pds5Δ elg1Δ cln2Δ* background and found that indeed *pds5Δ elg1Δ cln2Δ srs2Δ* strains are unable to lose the covering *PDS5* plasmid and are inviable on FOA plates.

Moreover, we could rescue this quadruple mutant by deleting the *RAD51* gene, encoding an ssDNA binding protein involved in homologous recombination, and substrate of Srs2 **(Figure 6C**). Therefore, in summary, we have found that *elg1Δ* promotes cohesion in the absence of Pds5 by accumulating SUMOylated PCNA on chromatin, thus promoting Srs2 activity to remove Rad51 filaments from ssDNA. We propose that by removing the Rad51 nucleoprotein, Srs2 generates ssDNA, which allows the deposition of cohesin molecules to establish sister chromatid cohesion when Pds5 is not present.

## DISCUSSION

Sister chromatid cohesion plays a fundamental role in cell division by ensuring faithful chromosome segregation. The establishment of sister chromatid cohesion is intimately linked to DNA replication, and many *bona fide* replication factors have been shown to be essential for cohesion establishment (13, 52, 53). In this study, we aimed to explore the genetic interactions between the PCNA unloader Elg1 and the cohesin accessory subunit Pds5. Although previous work showed that the deletion of *ELG1* could allow a temperature-sensitive *pds5-1* strain to grow at higher temperatures (22), the mechanistic details of this genetic interaction were not well understood.

Our genetic screens show that cells can retain SCC and viability in the absence of Pds5, if the essential functions provided by this protein are supplied by two alternative routes. We show that Pds5 protein is critical to protect cohesin function that is limited by Cln2-dependent inhibition of the *MCD1* transcription at the G1/S transition. We also show that the loss of cohesion caused by Pds5 deficiency can be partially suppressed by ectopic overexpression of *MCD1* or by deletion of *CLN2* **(Figure 1)**. Our results indicate that *cln2Δ* enhances cohesin function by promoting MBF activity, and thus *MCD1* cell cycle-dependent transcription at the G1/S transition. Thus the set point for cellular cohesin function is below its potential capacity because of limiting *MCD1* transcription early in the cell cycle. The notion that Mcd1 transcription limits cohesin function to suboptimal levels has precedent in recent studies of Ewing Sarcoma (54). These studies demonstrated that EWS-FLS1 fusion, a key determinant of this cancer, causes replicative stress and cellular senescence. The acquisition of an extra copy of the RAD21 (human ortholog of MCD1) dampens this stress and increases cell proliferation. Thus, also in these cells the level of Rad21 expression is suboptimal for addressing replicative stress (54). The existence of a suboptimal set point for MCD1 transcription for cohesion and DNA repair infers that optimal levels may have counteracting deleterious effects, for example inhibiting chromosome segregation or cohesin-independent pathways of DNA repair. Indeed, artificially limiting the Mcd1 levels by quantized reductions (QR) approach affect the Chromosome condensation, repetitive DNA stability, and DNA repair in yeast (55). While previous studies have not revealed phenotypes for cells overexpressing MCD1, our study suggests that a more comprehensive characterization of chromosome segregation, DNA repair and transcription in these cells is warranted.

Thus, our work helps delineate the molecular roles played by the Pds5 cohesin accessory factor.

### Pds5 is a cohesin stabilizer during S-phase

Cells lacking Pds5 protein exhibit high levels of premature separation of sister chromatids, which eventually jeopardize the chromosomal segregation program and result in cell death (**Figure 3)**; (20, 30, 35). Previous work showed that deletion of the SUMO E3 ligase Siz2 can rescue the temperature sensitivity and cohesion defects of the *pds5-1* temperature-sensitive strain by protecting the cohesin subunit Mcd1 from SUMO-dependent degradation (20). These results imply that Pds5 exerts a protective effect, and in its absence, Mcd1 is degraded, leading to the disintegration of cohesin complexes and to premature sister separation. However, overexpression of Mcd1 from high-copy number plasmids or by deleting the G1 cyclin *CLN2* was not sufficient to restore viability to cells completely lacking the Pds5 protein **(Figure 1)**. These results suggest that Pds5 plays several different roles in SCC. Our unbiased genetic screens help delineate them.

By using a degron allele of *PDS5*, we demonstrate that indeed, Mcd1 is quickly degraded following the auxin-induced degradation of Pds5, resulting in cell death. In contrast, we find that in the background of *elg1Δ cln2Δ,* Mcd1 protein no longer follows the sharp degradation kinetics associated with auxin-induced Pds5 degradation (**Figure 2**). Thus, decoupling the dependence of Mcd1 protein on Pds5 for its stability renders the *pds5Δ elg1Δ cln2Δ* strain viable. Altogether, our results show that Pds5 provides essential protection to the cohesin complex. Recently it is observed, that conditional degradation of Pds5 adversely affect the loop extrusion activity of a cohesin complex (56). The loop extrusion function of Pds5 is linked to its cohesin stabilization activity (57, 58). The observation that *pds5Δ elg1Δ cln2Δ* has sufficient cohesion (**Figure 3**) suggest that these cells stabilize cohesin complex in the absence of Pds5. In the future, it will be interesting to observe the cohesin’s loop extrusion activity in *elg1Δ cln2Δ* background.

### The G1 cyclin CLN2 as a novel suppressor of Pds5

In budding yeast, three G1 cyclins, *CLN1, CLN2,* and *CLN3,* are critical for starting the cell cycle and entry into subsequent cell cycle phases (59). These cyclins associate with the cell cycle-dependent kinase Cdc28 in a spatial and temporal manner to regulate the global gene expression. The Cln3 cyclin works upstream and is essential for the start of the cell cycle (60), where it activates the SBF and MBF transcription complexes. Cln1 and Cln2, on the other hand, are mainly involved in the G1/S transition and are believed to play functionally redundant roles (38).

We show that deletion of *CLN2,* but not *CLN1,* provides viability to a *pds5Δ elg1Δ* strain **(Figure 1)**. This result provides strong evidence that Cln1 and Cln2 are functionally distinct. The effect of *cln2Δ* is not due to increased stability of the Mcd1 protein, but rather to increased transcription of the *MCD1* gene by the MBF complex in the absence of *CLN2* (**Figure 4**). G1 cyclins Cln1 and Cln2 play a vital role in generating a phospho-degron on Sic1 protein, which is a potent S-phase inhibitor (61). The deletion of *CLN2* delays the entry into S-phase, prolonging the transcription period of *MCD1* and leading to an accumulation of its product. Thus, *cln2Δ*, similar to the high-copy number plasmid carrying the *MCD1* gene, rescues Pds5 deletion by providing an adequate amount of Mcd1 to compensate for its higher turnover in the absence of Pds5. These results establish an essential role of Pds5 in protecting Mcd1 at the G1/S boundary to insure proper SCC.

### ELG1 deletion promotes cohesion via SUMO-PCNA

In the absence of *ELG1,* cells accumulate PCNA on chromatin, both unmodified and SUMOylated (45). In the two-dot assays, the deletion of *ELG1* showed its effect mainly during SCC establishment and had only a minor effect during SCC maintenance **(Figure 3)**. By using different *elg1* alleles, we show that the ability of the different alleles to confer viability to a *pds5Δ cln2Δ* strain is negatively correlated with their sensitivity to DNA damaging agents (**Figure 5A**), reflecting their ability to unload PCNA from chromatin (46). Moreover, mutations in PCNA that lead to their spontaneous disassembly from chromatin (48) completely abolished the suppressive effect produced by deleting *ELG1*. Taken together, these results show that the suppression of *pds5Δ cln2Δ* is due to higher PCNA levels on the chromatin in the absence of the Elg1 PCNA unloader. The Eco1 acetyl-transferase binds PCNA, directly linking cohesion establishment to DNA replication (16). A simple model for the effect of deleting *ELG1* on the suppression of *pds5Δ* would therefore be through increased recruitment of the Eco1 acetyl-transferase. Unexpectedly, although high levels of PCNA on chromatin were observed in *elg1Δ,* the Eco1 levels on chromatin were not affected **(Figure 5C, D)**, ruling out this simple explanation. However, despite the lack of increase in Eco1 protein abundance at the fork, the level of Eco1-dependent Smc3 acetylation is elevated in *elg1Δ* mutants (62).

### *SUMOylated PCNA* recruits Srs2 to evict Rad51 from chromatin

Srs2 is a DNA helicase that evicts Rad51 filaments from the ssDNA and performs pro and anti-recombination roles during DNA replication (63, 64). Srs2 is recruited to chromatin by binding to SUMOylated PCNA (45), and has previously been shown to affect SCC (53). Our results show that Srs2 plays a central role in the pro-cohesion phenotype conferred by *elg1Δ*. Mutations that preclude SUMOylation of PCNA, or deletion of the *SRS2* gene itself, abolished the suppressive effect of *elg1Δ* and led to inviability of *pds5Δ elg1Δ cln2Δ* cells. Consistently with the known function of Srs2 function, the viability of a *pds5Δ elg1Δ cln2Δ srs2Δ* strain could be restored by deleting the *RAD51* gene, demonstrating that the role of *elg1Δ* is to recruit Srs2 in order to evict Rad51 from the chromatin **(Figure 6C)**.

What could be the consequence of Rad51 eviction? One possible explanation is that eviction of Rad51 exposes ssDNA and this is interpreted as a local DNA damage signal which may induce Eco1 activity and cohesion. This could be in principle the role played by Pds5 during S-phase. Importantly, this proposed mechanism is different from the known Chk1-dependent pathway in which DNA damage induces cohesion through acetylation of Mcd1 at lysines 84 and 210 (51) (**Figure 1D**). Similarly, a complete deletion of *CHK1* had no effect on the viability of a *pds5Δ elg1Δ cln2Δ* strain and did not prevent suppression of a *pds5Δ elg1Δ* strain by overexpression of Mcd1 (data not shown).

An alternative possibility is that Rad51 eviction allows the coupling between DNA replication and SCC establishment. Elegant biochemical assays by the Uhlmann’s lab recently established that cohesin can be loaded onto dsDNA, but second-strand entrapment requires ssDNA (65). They therefore suggested a model in which cohesin is loaded onto the dsDNA present on the leading strand at the moving fork, followed by entrapment of ssDNA at the lagging strand, which is then stabilized by further DNA synthesis (65). Thus, a stretch of protein-free ssDNA becomes essential for cohesion establishment. The ssDNA gaps left by Rad51’s eviction could thus allow more cohesion establishment in *elg1Δ.* Smc3 acetylation is a hallmark of stably established cohesion, and Smc3 acetylation protein levels are used as a proxy to monitor the extent of cohesion establishment during DNA replication (14). Consistent with our model, *elg1Δ* has a higher level of Smc3 acetylation than the wild type (62), suggesting that the absence of Elg1 promotes increased cohesion establishment, provided that an ample enough amount of Mcd1 protein is available.

### A model for the roles of Pds5 and the suppression of pds5Δ by elg1Δ cln2Δ

Our results delineate two essential roles for Pds5 in SCC: it protects the integrity of cohesin by preventing Mcd1 degradation, and it is involved in the activation of Smc3 acetylation by Eco1. These two roles take place during S-phase, and coordinate DNA replication with SCC.

Pds5 is necessary in order to protect the Mcd1 protein from SUMOylation and STUbL-dependent degradation (20, 21, 66). Deletion of both *CLN2* and *ELG1,* or overexpression of *MCD1* from a plasmid, contributes to increase Mcd1 levels. Whereas the first deletion increases MBF-dependent transcription of the *MCD1* gene (**Figure 4**), *ELG1* deletion may indirectly ensure higher levels of cohesive cohesin, in which, after Eco1 activity, Mcd1 may become resistant to degradation. However, the increase in the Mcd1 protein level is not sufficient to provide SCC in the absence of Pds5 (**Figure 7**). The second role for Pds5 occurs during DNA replication and involves the activation of Eco1 activity, required for stabilizing cohesin on the chromatin. This second activity can be supplied by a deletion of *ELG1,* provided enough Mcd1 is present. As we have shown, increased SUMO-PCNA on the chromatin allows increased cohesin loading and establishment by recruiting the Srs2 helicase to evict Rad51 (**Figure 6**). The increased SCC establishment explains the ability of *elg1Δ* to rescue the temperature sensitivity of both *pds5-1* and *eco1-1* strains (67)’(22), and is consistent with higher Smc3 acetylation levels (62) of *elg1Δ* mutants. Just increasing the rate of establishment, however, is not enough, if the level of Mcd1 is kept low due to its de-protection by the absence of Pds5. Only a combination of higher Mcd1 levels (provided by *cln2Δ* or by *MCD1* overexpression), together with the increased Rad51 eviction (indirectly caused by *ELG1* deletion) ensure a robust SCC in the total absence of Pds5 (**Figure 7**). In summary, our results thus provide novel insights on the function of the accessory cohesin subunit Pds5 in SCC.

**Figure 7.**
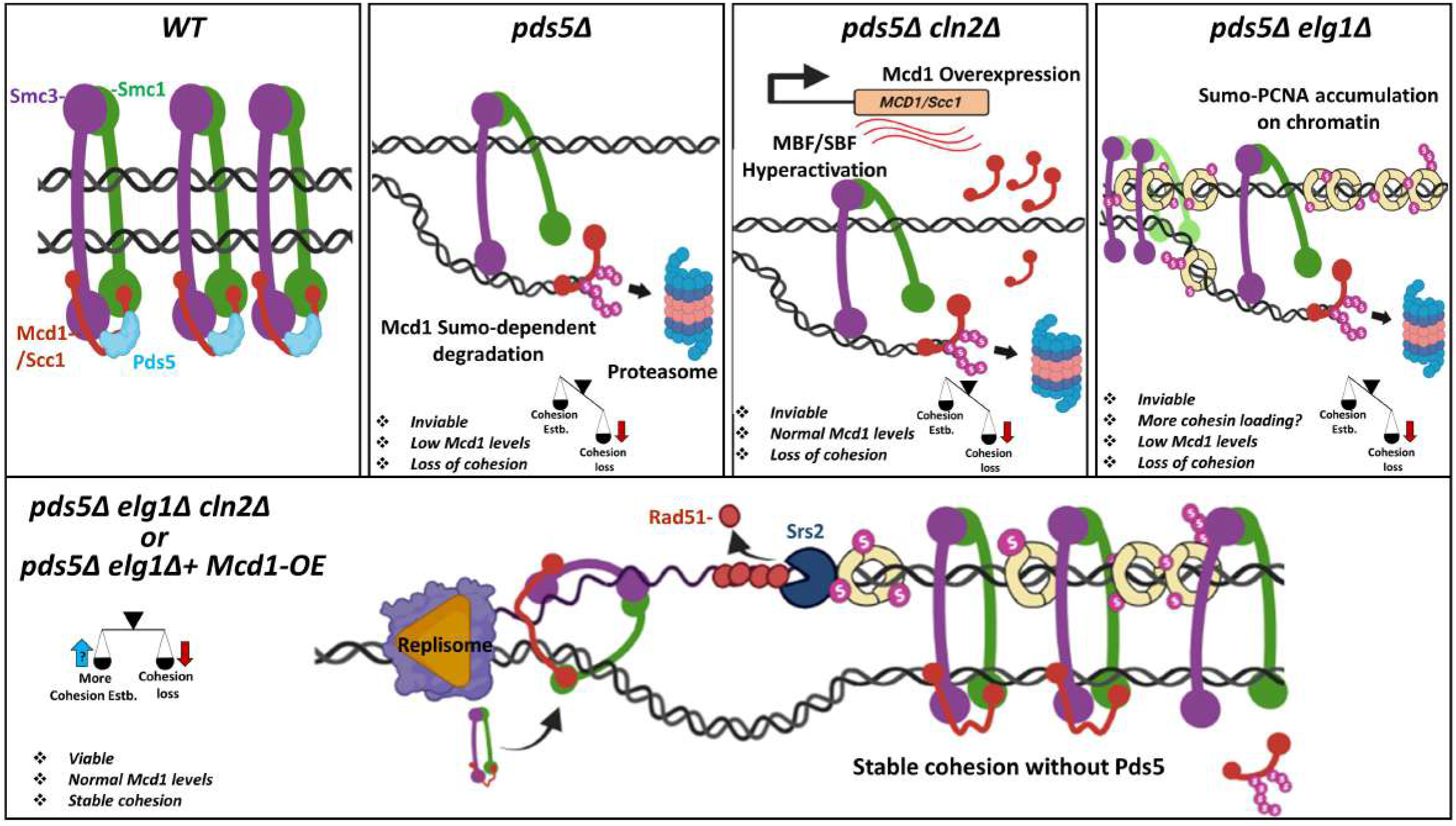
A model for the bypass of Pds5 function by *elg1*Δ *cln2*Δ. **A)** The Wt cells properly establish cohesion during the S-phase and maintain it throughout the following cell cycle to allow faithful chromosome segregation. **B)** The deletion of Pds5 results in hyper-SUMOylation of the Mcd1 cohesin subunit, leading to its premature degradation, followed by loss of cohesion and cell death. **C)** The deletion of the G1 cyclin Cln2 results in overproduction of Mcd1, however it cannot produce sufficient cohesion to sustain the high cohesin turnover associated with the loss of Pds5 protein. As a result, the *pds5*Δ *cln2*Δ strain is inviable and show cohesion defects. **D)** The deletion of PCNA unloader Elg1 results in accumulation of SUMO-PCNA on chromatin which might allow a wider window for cohesin establishment. However, *pds5*Δ *elg1*Δ strain is inviable due to the insufficient levels of Mcd1 protein available during cohesion establishment. **E)** The deletion of PCNA unloader Elg1 along with G1 cyclin Cln2 (or with Mcd1 over-expression) results in stable cohesion in the absence of the Pds5 cohesin subunit, rendering yeast cells viable. In other words, the high cohesin turnover associate with *pds5Δ* might be compensated by over establishing functional cohesion during DNA replication in this scenario. The SUMO-PCNA accumulation recruits Srs2 to remove Rad51 protein from ssDNA, which might allow the increased establishment of cohesion during DNA replication. (Estb. Stands for establishment).

## MATERIALS AND METHODS

### Yeast strains and media

All yeast strains used in this study are of A364A background. YPD medium was prepared with a ready-to-use mixture (FORMEDIUM). SC minimal was prepared with 2% dextrose (FORMEDIUM), Yeast Nitrogen Base w/o Amino Acids (DIFCO), and all necessary amino acids. 2% of agar (DIFCO) was added for solid media. Auxin (3-indole acetic acid; Sigma-Aldrich Catalogue # I3705) was added to SC minimal media with 300 uM final concentration in DMSO. 5-FOA is SD with all amino acids and nucleobases, but only 50 mg of uracil and 0.8 g of 5-fluoroortic acid (5-FOA) were used per liter of media.

### Cell cycle arrest

For experiments requiring cell cycle arrest, cells were grown at 30°C in SC complete medium until mid-log phase (0.6 OD_600_) and incubated with nocodazole (Sigma-Aldrich; Catalogue # M1404) (15μg/ml) for G2-M arrest or alpha-factor (Sigma-Aldrich; Catalogue # T6901) (50 ng/mL) for G1 arrest. Both incubation times were of two-hour duration. The text figures legends mention all cell cycle arrest experiment details.

### Yeast spot assays

Cells were grown to saturation in SC media at 30°C, diluted to 1 OD_600_, and then plated in 5-fold serial dilutions. Cells were incubated on plates at 30°C for 3-5 days. 10 μL from each appropriate dilution were then spotted on respective plates.

### Yeast genetic screen for the suppressors of *pds5Δ elg1Δ*

For the high copy number suppressor screen, the yeast cells were transformed with the entire Prelich collection, consisting of over 1500 plasmids containing a unique clone of a segment of the yeast *S. cerevisiae* genome. The cells were plated on 5-FOA plates to lose the Pds5 covering plasmid. The colonies that grew on 5-FOA were confirmed for the loss of covering plasmid followed by Plasmids isolation and sequencing. The library was constructed by partially digesting prototrophic yeast genomic DNA with MboI and subcloning it into the BamHI sites of the *E. coli*-yeast shuttle vector, pGP564. The proteins are untagged and expressed from their endogenous wild-type promoter. The pGP564 shuttle vector contains the *LEU2* selectable marker and 2-micron plasmid sequences necessary to maintain a high copy number in yeast. The average insert size in this library is approximately 10 kb, with each insert containing an average of 4-5 genes. **B)** For the spontaneous suppressor screen, the cells carrying a double deletion of *PDS5* and *ELG1* and a *URA3-PDS5-LEU2* covering plasmid were plated on 5-FOA plates. Cells that grew on 5-FOA and were also Leu- (i.e., lost the covering plasmid) were subjected to whole-genome sequencing to find suppressor mutations in the genome.

### Whole-genome sequencing of yeast strains

Sequencing libraries were constructed for each strain from whole-genome DNA, using a small-volume Nextera (Illumina.com
) tagmentation protocol (68). Unique combinations of Nextera dual-index adapters were used for each sample, and all samples were multiplexed onto one Illumina HiSeq 2000 lane. Sequencing was performed at the Stanford Center for Genomics and Personalized Medicine using 2×101bp paired-end read technology. Variant calling was carried out using CLC Genomics Workbench v8.5 (Qiagen.com). Sequences were uploaded to the NIH SRA under project number **PRJNA742489**.

### Cohesion analysis using the LacO-LacI system

We monitored the cohesion establishment and maintenance using the LacO-LacI system. Briefly, cells carrying tandem LacO repeats integrated at *LYS4,* located 470 kb from *CEN4,* and a GFP-LacI fusion was used. For establishment experiments, cells were grown at 30°C in SC minimal medium until mid-log phase (0.6 OD_600_) and then incubated with alpha-factor (50 ng/mL) for G1 arrest for 2 hours. For depletion of AID-Pds5, Auxin was added (300 uM) simultaneously. After this incubation, cells were washed three times in YPD (30°C) containing 0.1 mg/ml Pronase E (Sigma-Aldrich; Catalogue # P5147), resuspended in SC minimal medium containing nocodazole (15μg/ml), and then incubated at 30°C for 2 h to early mitosis arrest while cohesion disjunction was analyzed every 20 min. For maintenance experiments, cells were grown at 30°C in SC minimal medium until mid-log phase (0.6 OD_600_) and then incubated with nocodazole (15μg/ml) for 2 hours. After this incubation, auxin was added (300 uM) for the depletion of AID-Pds5 proteins together with nocodazole (15μg/ml) for 2 h at 30°C while cohesion disjunction was analyzed every 20 min. Images were acquired with an EVO FL microscope (ThermoFisher Scientific; Catalogue #AMF4300) equipped with the GFP Light Cube (470/22 nm Excitation; 510/42 nm Emission) (ThermoFisher Scientific; Catalogue #AMEP4651).

### Flow Cytometry

For yeast cell cycle examination using Flow cytometry, the protocol by Harari et al. 2018 (69) was used. Briefly, For a given time point, cells were spun down, washed with 200 μL TE solution (10mM Tris-HCl pH 7.5, 1mM EDTA), resuspended in 60 μL of TE, and fixated by adding 140 μL of absolute cold ethanol and incubated overnight at 4°C. Cells were then washed twice using TE buffer, resuspended in 100 μL of TE-RNase solution (10mM Tris-HCl pH 7.5, 1mM EDTA, and 0.25mg/mL RNase) incubated for 2 h at 37°C. Cells were then rewashed using TE buffer, resuspended in 200 μL of proteinase-K solution (10mM Tris-HCl pH 7.5, 1mM EDTA, and 0.25mg/mL proteinase-K) incubated for 2 h at 37°C. Cells were then again washed using TE buffer and resuspended in 200 μL of TE-PI buffer (Tris EDTA and 20 μg/mL Propidium-iodide) and incubated overnight at 4°C in the dark. Before measuring, samples were sonicated three times for 2s at 20% intensity and checked under the microscope for the absence of cell clusters/doublets. All samples were analyzed using a Flow cytometry MACSQuant system, and Flow data were analyzed using FlowJo programs. Doublets were eliminated using a pulse geometry gate (FSC-H x FSC-A). In order to measure the mean fluorescent intensity, yeast cells carrying the GFP/mCherry plasmids were harvested in the mid-log phase (O.D_600_ ~ 0.6) and washed twice with TE buffer (10mM Tris-HCl pH 7.5, 1mM EDTA) and subjected to flow cytometer after resuspending in TE buffer. Around 25000 events were monitored, and samples were analyzed using the FlowJo program. The events were aligned on the ds-Red_txRed-H channel for mCherry and GFP_FITC-H for eGFP. Five independent (n=5) replicates were performed for all samples.

### Chromatin Fractionation

The protocol used for chromatin enrichment is described in (70). Around 40 OD cells were harvested from a logarithmically growing yeast culture and resuspended in 1 mL of pre-spheroplasting buffer (100 mM PIPES/KOH, pH 9.4, 10 mM DTT, 0.1% sodium azide). Cells were transferred to 1.5 ml tubes and incubated on ice for 10min with a brief vortex in between. Next, cells were suspended in spheroplasting buffer (50 mM KH_2_PO_4_/K_2_HPO_4_, pH 7.4, 0.8 M sorbitol, 10 mM DTT, 0.1% sodium azide) containing 200 μg/ml Zymolyase-100T at 30°C for 30 mins on a roller at slow speed. The spheroplasts were confirmed microscopically, and protocol from (70) was followed afterward. The Histone H3 and Rps6 were used as a control for chromatin enrichment.

### Protein extraction, Western blotting, antibodies, and band quantitation

Cells equivalents of 3 OD_600_ were pelleted and stored at −80°C. Proteins were extracted from cells as described previously (71) using either a tri-chloroacetic acid method(72). To resolve Pds5, Mcd1, and Tubulin, 8% SDS-polyacrylamide gels were used. Immunoblotting was done as described previously. To detect proteins, the following primary antibodies were used: Anti-Mcd1 (1:10000), Anti-sV5 Santa Cruz (sc-58052) (1:1000), Anti-Actin Abcam (Ab8226)1:1000, Anti-tubulin (1:1000), Anti-GFP Abcam (Ab290) 1:1000. Anti-H3 (ab1791) Abcam 1:1000, Anti-RPS6 (ab40820) Abcam 1:1000, Anti-PCNA (ab70472) Abcam 1:1000 Anti-MYC (9E10, SC-40) Santa Cruz 1:1000 and Anti-HA (sc7392) Santa Cruz 1:1000. Western blot bands were quantified with ImageJ (www.imagej.net).

## ACKNOWLEDGMENT

We thank Doug Koshland for support, ideas and comments on the paper. We thank all members of the Kupiec lab and Koshland lab for helpful discussions and encouragement. Work in the Kupiec lab was supported by grants from the Israel Science Foundation, the Israel Cancer Research Fund and the Minerva Stiftung.

## COMPETING INTEREST STATEMENT

None declared

## SUPPLEMENTARY FIGURE LEGENDS

**Figure S1. Screen for the suppressors of *pds5Δ elg1Δ.* A)** Spot assay with fivefold serial dilutions of the *pds5Δ* and *pds5Δ elg1Δ* strains carrying Pds5 centromeric *URA* covering plasmid on SD-Ura and 5-FOA plates. **B)** Anti-Mcd1 western blot shows the overexpression of different mcd1 mutants in pds5Δ and *pds5Δ elg1Δ* strains compared to empty vector. Actin (probed with anti-Actin Ab) was used as a loading control. The graph below the western blot panel represent the average (n=3, mean ± SD) fold change in the Mcd1 expression levels compared to the empty vector. **C)** List of *de novo* mutations observed in G1 cyclin *CLN2* gene that allow the *pds5Δ elg1Δ* strain viability.

**Figure S2. Auxin induced degradation of *AID-PDS5*. A)** Western blot showing the degradation kinetics of Pds5 protein on addition of Auxin (IAA, 300 μM) to the growth media. **B)** Quantification of the Pds5 protein levels at the indicated time point normalized to the tubulin loading control (% mean ± SD; n=3). T test analysis p value ≤ 0.01. **C)** Flow cytometry data supporting the *Figure no. 2 A, B.* Data represents that the cells were arrested in G2/M phase while they were harvested for protein extraction at the final time point T120min.

**Figure S3. Mcd1 protein half-life unchanged in *elg1Δ cln2Δ* strain. A)** Western blot for the cycloheximide chase experiments in *PDS5-AID* and *PDS5-AID elg1Δ cln2Δ* strain. The cells were grown until log phase (time 0) followed by treatment with cycloheximide CHX (250 μg/mL). Samples were taken every 20 minutes until completing a 2 hours experiment. **B)** Quantification of the Mcd1 and Pds5 protein levels normalized to tubulin as loading control (n=3; % mean ± SD). **C)** Statistical analysis of the Mcd1 half-life by performing T test analysis (ns = not significant). **D)** Western blot for the Auxin+ cycloheximide chase experiments in *PDS5-AID* and *PDS5-AID elg1Δ cln2Δ* strain. The cells were grown until log phase (time 0) followed by treatment with Auxin (IAA 300 μM) along with cycloheximide CHX (250 μg/mL). Samples were taken every 20 minutes until completing a 2 hours experiment. **E)** Quantification of the Mcd1 and Pds5 protein levels normalized to tubulin as loading control (n=3; % mean ± SD). **F)** Statistical analysis of the Mcd1 half-life by performing T test analysis (ns = not significant).

---

| YEAST-Strain number | Genotype |
| --- | --- |
| **MKDK23** | *Mat A pds5Δ::Hygmx lacO (DK)-NAT; 10kbCEN4 pHIS3-GFPLacI-HIS3::his3-11,15 GAL1+ trp1-1 leu2-3,112 ura3-52 bar1 + pGV282 [CEN3 URA3 PDS5]* |
| **MKDK113** | *Mat A pds5Δ::Hygmx elg1Δ :: KanMX lacO (DK)-NAT; 10kbCEN4 pHIS3-GFPLacI-HIS3::his3-11,15 GAL1+ trp1-1 leu2-3,112 ura3-52 bar1 + pGV282 [CEN3 URA3 PDS5]* |
| **MKDK470** | *Mat A pds5Δ::Hygmx cln2Δ :: cgTRP1 lacO (DK)-NAT; 10kbCEN4 pHIS3-GFPLacI-HIS3::his3-11,15 GAL1+ trp1-1 leu2-3,112 ura3-52 bar1 + pGV282 [CEN3 URA3 PDS5]* |
| **MKDK471** | *Mat A pds5Δ::Hygmx elg1Δ :: KanMX cln2Δ :: cgTRP1 lacO (DK)-NAT; 10kbCEN4 pHIS3-GFPLacI-HIS3::his3-11,15 GAL1+ trp1-1 leu2-3,112 ura3-52 bar1 + pGV282 [CEN3 URA3 PDS5]* |
| **MKDK474** | *Mat A pds5Δ::Hygmx cln1Δ :: cgTRP1 lacO (DK)-NAT; 10kbCEN4 pHIS3-GFPLacI-HIS3::his3-11,15 GAL1+ trp1-1 leu2-3,112 ura3-52 bar1 + pGV282 [CEN3 URA3 PDS5]* |
| **MKDK477** | *Mat A pds5Δ::Hygmx elg1Δ :: KanMX cln1Δ :: CgTRP1 lacO (DK)-NAT; 10kbCEN4 pHIS3-GFPLacI-HIS3::his3-11,15 GAL1+ trp1-1 leu2-3,112 ura3-52 bar1 + pGV282 [CEN3 URA3 PDS5]* |
| **MKDK38** | *Mat A LacO-NAT::lys4 trp1-1 bar1 GFPLacI-HIS3::his3-11,15 leu2-3,112 ura3-52 GAL+* |
| **MKDK475** | *Mat A PDS5-3v5-AID2::KanMX6 ADH1-TIR1-URA3::ura3-52 LacO(DK)-NAT::lys4 pHIS3-GFP-LacI-HIS3::his3-11,15 trp1-1 leu2-3,112 bar1 GAL+* |
| **E-B1-62** | *Mat A PDS5-3v5-AID2::KanMX6 elg1Δ::HygMX ADH1-TIR1-URA3::ura3-52 LacO(DK)-NAT::lys4 pHIS3-GFP-LacI-HIS3::his3-11,15 trp1-1 leu2-3,112 bar1 GAL+* |
| **E-B1-64** | *Mat A PDS5-3v5-AID2::KanMX6 cln2Δ::cgTRP1 ADH1-TIR1-URA3::ura3-52 LacO(DK)-NAT::lys4 pHIS3-GFP-LacI-HIS3::his3-11,15 trp1-1 leu2-3,112 bar1 GAL+* |
| **E-B1-73** | *Mat A PDS5-3v5-AID2::KanMX6 elg1Δ::HygMX cln2Δ::cgTRP1 ADH1-TIR1-URA3::ura3-52 LacO(DK)-NAT::lys4 pHIS3-GFP-LacI-HIS3::his3-11,15 trp1-1 leu2-3,112 bar1 GAL+* |
| **SC_190** | *Mat A Pds5-3v5-AID2::KanMX6 ADH1-TIR1-URA3::ura3-52 his3-11,15 trp1-1 leu2-3,112 lys2-801, bar1 GAL+* |
| **SC_193** | *Mat A Pds5-3v5-AID2::KanMX elg1Δ::HygMX ADH1-TIR1-URA3::ura3-52 his3-11,15 trp1-1 leu2-3,112 lys2-801, bar1 GAL+* |
| **SC_196** | *Mat A Pds5-3v5-AID2::KanMX cln2Δ::cgTRP1 ADH1-TIR1-URA3::ura3-52 his3-11,15 trp1-1 leu2-3,112 lys2-801, bar1 GAL+* |
| **SC_199** | *Mat A Pds5-3v5-AID2::KanMX elg1Δ::HygMX cln2Δ::cgTRP1 ADH1-TIR1-URA3::ura3-52 his3-11,15 trp1-1 leu2-3,112 lys2-801, bar1 GAL+* |
| **SC_267** | *Mat A Elg1(WT)-13myc::KanMX pds5Δ::Hygmx cln2Δ :: cgTRP1 lacO (DK)-NAT; 10kbCEN4 pHIS3-GFPLacI-HIS3::his3-11,15 GAL1+ trp1-1 leu2-3,112 ura3-52 bar1 +CEN PDS5 URA* |
| **SC_268** | *Mat A 3XSIM-ELG1 -13myc::KanMX pds5Δ::Hygmx cln2Δ :: cgTRP1 lacO (DK)-NAT; 10kbCEN4 pHIS3-GFPLacI-HIS3::his3-11,15 GAL1+ trp1-1 leu2-3,112 ura3-52 bar1 +CEN PDS5 URA* |
| **SC_269** | *Mat A elg1-386/7AA-13MYC::KanMX pds5Δ::Hygmx cln2Δ :: cgTRP1 lacO (DK)-NAT; 10kbCEN4 pHIS3-GFPLacI-HIS3::his3-11,15 GAL1+ trp1-1 leu2-3,112 ura3-52 bar1 +CEN PDS5 URA* |
| **SC_270** | *Mat A elg1-386/7DD-13MYC::KanMX pds5Δ::Hygmx cln2Δ :: cgTRP1 lacO (DK)-NAT; 10kbCEN4 pHIS3-GFPLacI-HIS3::his3-11,15 GAL1+ trp1-1 leu2-3,112 ura3-52 bar1 +CEN PDS5 URA* |
| **SC_271** | *Mat A 3X-SIM + elg1-386/7DD-13MYC::KanMX pds5Δ::Hygmx cln2Δ :: cgTRP1 lacO (DK)-NAT; 10kbCEN4 pHIS3-GFPLacI-HIS3::his3-11,15 GAL1+ trp1-1 leu2-3,112 ura3-52 bar1 +CEN PDS5 URA* |
| **SC_272** | *Mat A elg1-KK343/4AA-13myc::KanMX pds5Δ::Hygmx cln2Δ :: cgTRP1 lacO (DK)-NAT; 10kbCEN4 pHIS3-GFPLacI-HIS3::his3-11,15 GAL1+ trp1-1 leu2-3,112 ura3-52 bar1 +CEN PDS5 URA* |
| **SC_273** | *Mat A elg1-KK343/4DD-13myc::KanMX pds5Δ::Hygmx cln2Δ :: cgTRP1 lacO (DK)-NAT; 10kbCEN4 pHIS3-GFPLacI-HIS3::his3-11,15 GAL1+ trp1-1 leu2-3,112 ura3-52 bar1 +CEN PDS5 URA* |
| **SC_274** | *Mat A elg1-DD407,409AA-13myc::KanMX pds5Δ::Hygmx cln2Δ :: cgTRP1 lacO (DK)-NAT; 10kbCEN4 pHIS3-GFPLacI-HIS3::his3-11,15 GAL1+ trp1-1 leu2-3,112 ura3-52 bar1 +CEN PDS5 URA* |

| Strain number | Genotype |
| --- | --- |
| **SC_275** | *Mat A elg1-DD407,409AA + KK343/344AA -13myc::KanMX pds5Δ::Hygmx cln2Δ :: cgTRP1 lacO (DK)-NAT; 10kbCEN4 pHis3-GFP<sub>LacI</sub>-His3:his3-11,15 GAL1+ trp1-1 leu2-3,112 ura3-52 bar1 +CEN PDS5 URA* |
| **SC_276** | *Mat A elg1-DD407,409AA+ KK343/344DD-13myc::KanMX pds5Δ::Hygmx cln2Δ :: cgTRP1 lacO (DK)-NAT; 10kbCEN4 pHis3-GFP<sub>LacI</sub>-His3:his3-11,15 GAL1+ trp1-1 leu2-3,112 ura3-52 bar1 +CEN PDS5 URA* |
| **SC_277** | *Mat A 3X-SIM + elg1-386/7AA-13MYC::KanMX pds5Δ::Hygmx cln2Δ :: cgTRP1 lacO (DK)-NAT; 10kbCEN4 pHis3-GFP<sub>LacI</sub>-His3:his3-11,15 GAL1+ trp1-1 leu2-3,112 ura3-52 bar1 +CEN PDS5 URA* |
| **SC_99** | *Mat A pds5Δ::Hygmx leu2::pol30-D150E elg1Δ :: KanMX cln2Δ :: cgTRP1 lacO (DK)-NAT; 10kbCEN4 pHis3-GFP<sub>LacI</sub>-His3:his3-11,15 GAL1+ trp1-1 leu2-3,112 ura3-52 bar1 + pGV282 [CEN3 URA3 PDS5]* |
| **SC_100** | *Mat A pds5Δ::Hygmx leu2::pol30-E143K elg1Δ :: KanMX cln2Δ :: cgTRP1 lacO (DK)-NAT; 10kbCEN4 pHis3-GFP<sub>LacI</sub>-His3:his3-11,15 GAL1+ trp1-1 leu2-3,112 ura3-52 bar1 + pGV282 [CEN3 URA3 PDS5]* |
| **SC_93** | *Mat A pds5Δ::Hygmx leu2::pol30-S152P elg1Δ :: KanMX cln2Δ :: cgTRP1 lacO (DK)-NAT; 10kbCEN4 pHis3-GFP<sub>LacI</sub>-His3:his3-11,15 GAL1+ trp1-1 leu2-3,112 ura3-52 bar1 + pGV282 [CEN3 URA3 PDS5]* |
| **SC_310** | *Mat A Eco1-3HA::HisMx6 Pds5-3v5-AID2::KanMX ADH1-TIR1-URA3::ura3-52 his3-11,15 trp1-1 leu2-3,112 lys2-801, bar1 GAL+* |
| **SC_311** | *Mat A Eco1-3HA::HisMx6 elg1 Δ::HygMX Pds5-3v5-AID2::KanMX ADH1-TIR1-URA3::ura3-52 his3-11,15 trp1-1 leu2-3,112 lys2-801, bar1 GAL+* |
| **SC_73** | *Mat A pds5Δ::Hygmx leu2::pol30 K127R elg1Δ :: KanMX cln2Δ :: cgTRP1 lacO (DK)-NAT; 10kbCEN4 pHis3-GFP<sub>LacI</sub>-His3:his3-11,15 GAL1+ trp1-1 leu2-3,112 ura3-52 bar1 + pGV282 [CEN3 URA3 PDS5]* |
| **SC_74** | *Mat A pds5Δ::Hygmx leu2::pol30 K127R,K164R elg1Δ :: KanMX cln2Δ :: cgTRP1 lacO (DK)-NAT; 10kbCEN4 pHis3-GFP<sub>LacI</sub>-His3:his3-11,15 GAL1+ trp1-1 leu2-3,112 ura3-52 bar1 + pGV282 [CEN3 URA3 PDS5]* |
| **SC_75** | *Mat A pds5Δ::Hygmx leu2::pol30 K164R elg1Δ :: KanMX cln2Δ :: cgTRP1 lacO (DK)-NAT; 10kbCEN4 pHis3-GFP<sub>LacI</sub>-His3:his3-11,15 GAL1+ trp1-1 leu2-3,112 ura3-52 bar1 + pGV282 [CEN3 URA3 PDS5]* |
| **SC_108** | *Mat A rad5Δ::KanMX elg1Δ::LEU2-MX pds5Δ::Hygmx cln2Δ :: cgTRP1 lacO (DK)-NAT; 10kbCEN4 pHis3-GFP<sub>LacI</sub>-His3:his3-11,15 GAL1+ trp1-1 leu2-3,112 ura3-52 bar1 + pGV282 [CEN3 URA3 PDS5]* |
| **SC_159** | *Mat A rad18Δ::KanMX elg1Δ :: HisGMX pds5Δ::Hygmx cln2Δ :: cgTRP1 lacO (DK)-NAT; 10kbCEN4 pHis3-GFP<sub>LacI</sub>-His3:his3-11,15 GAL1+ trp1-1 leu2-3,112 ura3-52 bar1+ pGV282 [CEN3 URA3 PDS5]* |
| **SC_110** | *Mat A siz1Δ::KanMX elg1Δ::LEU2-MX pds5Δ::Hygmx cln2Δ :: cgTRP1 lacO (DK)-NAT; 10kbCEN4 pHis3-GFP<sub>LacI</sub>-His3:his3-11,15 GAL1+ trp1-1 leu2-3,112 ura3-52 bar1 + pGV282 [CEN3 URA3 PDS5]* |
| **SC_111** | *Mat A srs2Δ::KanMX elg1Δ::LEU2-MX pds5Δ::Hygmx cln2Δ:: cgTRP1 lacO (DK)-NAT; 10kbCEN4 pHis3-GFP<sub>LacI</sub>-His3:his3-11,15 GAL1+ trp1-1 leu2-3,112 ura3-52 bar1 + pGV282 [CEN3 URA3 PDS5]* |
| **SC_266** | *Mat A rad51Δ::Leu2 srs2Δ::KanMX elg1Δ :: HisGMX pds5Δ::Hygmx cln2Δ :: cgTRP1 lacO (DK)-NAT; 10kbCEN4 pHis3-GFP<sub>LacI</sub>-His3:his3-11,15 GAL1+ trp1-1 leu2-3,112 ura3-52 bar1 + pGV282 [CEN3 URA3 PDS5]* |
| **SX48B** | *Mat A trp1-1::TIR1-cgTRP1 LacO-NAT::lys4 GFP<sub>LacI</sub>-His3:his3-11,15 SMC3-A1089-V5-BirA SMC3-P533-Avi6HA-LEU2:leu2-3,112 ura3-52 bar1 GAL+* |
| **SX122B** | *Mat A trp1-1::TIR1-cgTRP1 LacO-NAT::lys4 GFP<sub>LacI</sub>-His3:his3-11,15 SMC3-A1089-V5-BirA SMC3-P533-Avi6HA-LEU2:leu2-3,112 PDS5-3V5-AID2:G418 ura3-52 bar1 GAL+* |
| **SX283** | *Mat A trp1-1::TIR1-cgTRP1 LacO-NAT::lys4 GFP<sub>LacI</sub>-His3:his3-11,15 SMC3-A1089-V5-BirA SMC3-P533-Avi6HA-LEU2:leu2-3,112 cln2Δ::cgTRP1 elg1Δ::HygMX ura3-52 bar1 GAL+* |
| **SX284** | *Mat A trp1-1::TIR1-cgTRP1 LacO-NAT::lys4 GFPLacI-HIS3:his3-11,15 SMC3-A1089-V5-BirA SMC3-P533-Avi6HA-LEU2:leu2-3,112 cln2Δ::cgTRP1 elg1Δ::HygMX PDS5-3V5-AID2:G418 ura3-52 bar1 GAL+* |
| **SX297** | *Mat A trp1-1::TIR1-cgTRP1 LacO-NAT::lys4 GFPLacI-HIS3:his3-11,15 SMC3-A1089-V5-BirA SMC3-P533-Avi6HA-LEU2:leu2-3,112 elg1Δ::HygMX PDS5-3V5-AID2:G418 ura3-52 bar1 GAL+* |
| **SX298** | *Mat A trp1-1::TIR1-cgTRP1 LacO-NAT::lys4 GFPLacI-HIS3:his3-11,15 SMC3-A1089-V5-BirA SMC3-P533-Avi6HA-LEU2:leu2-3,112 cln2Δ::cgTRP1 PDS5-3V5-AID2:G418 ura3-52 bar1 GAL+* |

## PLASMIDS

PLASMIDS:
| Plasmid number | Insert information |
| --- | --- |
| **pGV282** | *CEN3 URA3 pPds5-PDS5* |
| **MKDK400** | *YEp181-2μ-LEU2 pMcd1-MCD1 (WT)* |
| **MKDK402** | *YEp181-2μ-LEU2 pMcd1-mcd1-KK84,210QQ* |
| **MKDK404** | *YEp181-2μ-LEU2 pMcd1- mcd1-KK84,210RR* |
| **MKDK327** | *YEp181-2μ-LEU2 pMcd1- mcd1-F528R* |
| **MKDK329** | *YEp181-2μ-LEU2 pMcd1-mcd1-L532R* |
| **MKDK335** | *YEp181-2μ-LEU2 pMcd1-mcd1-V137K* |
| **K133** | *pRS425-2μ-LEU2 pADH1-mCherry pMcd1-yEGFP-CL1 (degron)* |
| **K177** | *pRS425-2μ-LEU2 pADH1-mCherry pMcd1 Δ(-372 to -366) -yEGFP-CL1 (degron) [ΔMCB-DISTAL]* |
| **K179** | *pRS425-2μ-LEU2 pADH1-mCherry pMcd1 Δ(-292 & -286) -yEGFP-CL1 (degron) [ΔMCB-PROXIMAL]* |

## Notes

### Competing Interest Statement

The authors have declared no competing interest.

