## Supplementary material for "*S. cerevisiae* cells can grow without the Pds5 cohesin subunit": Suppl. Figure 1

**A**

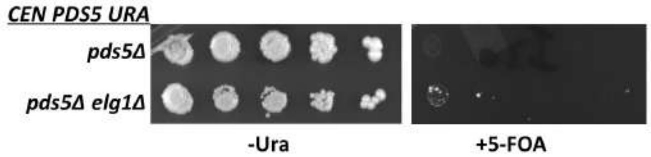


**B**

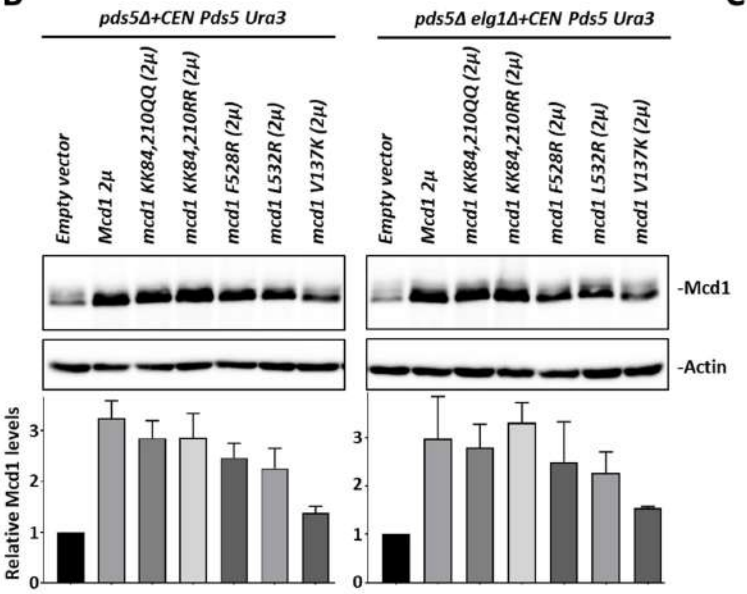


**C**

| GENE NAME | Mutation | No. of times mutation appeared |
|---|---|---|
| CLN2 | Ser 315 (* stop codon) | 2x |
| CLN2 | Arg 100 Ile | 2x |
| CLN2 | Gly132 fs (frame shift) | 4x |
| CLN2 | Ile186 fs (frame shift) | 2x |
| CLN2 | Lys 225 fs (frame shift) | 2x |
| CLN2 | Asp 273 (gene deletion) | 3x |
| CLN2 | Ile 94 leu, His 97 fs (frame shift) | 1x |
| CLN2 | Arg 9 fs (frame shift) | 2x |
| CLN2 | Gly 132 fs (frame shift) | 3x |
| CLN2 | Ty insertion | 1x |
| CLN2 | Pro 470 fs (frame shift) | 1x |

**Figure S1**
