## Supplementary figures and images for "*S. cerevisiae* cells can grow without the Pds5 cohesin subunit"

### Suppl. Figure 2

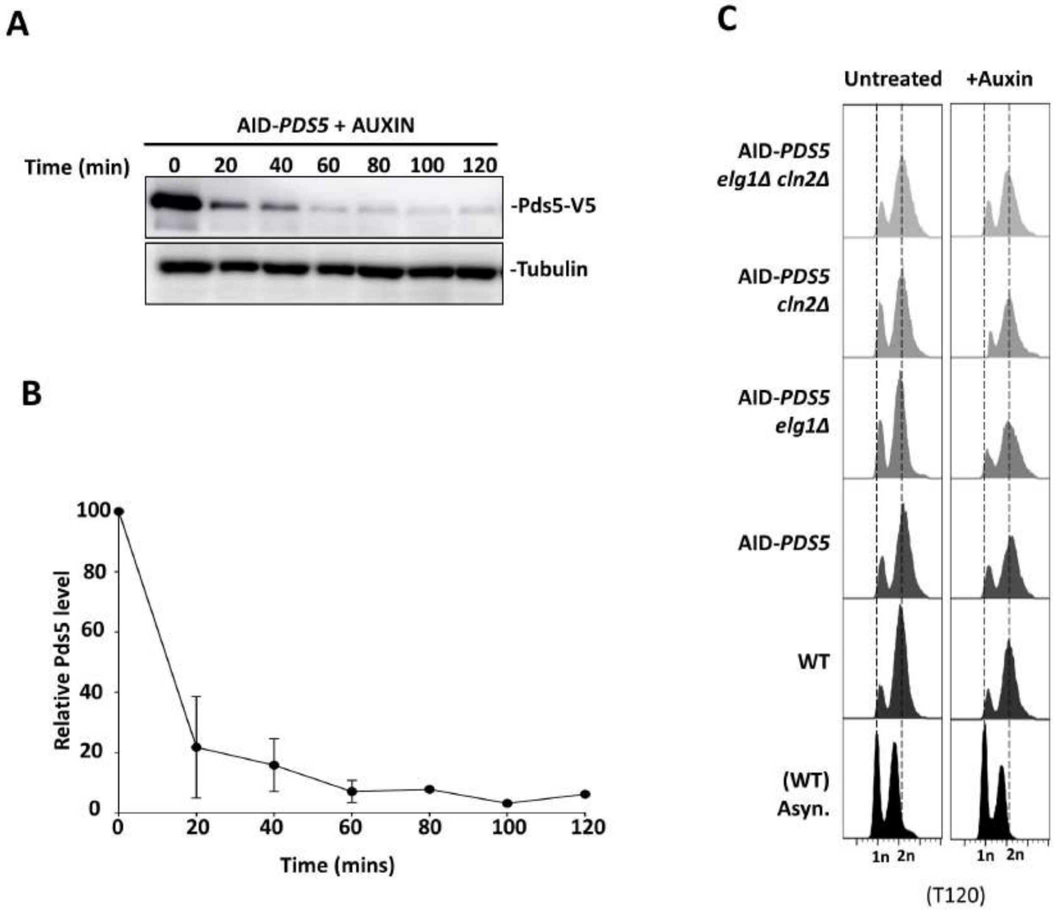

### Suppl. Figure 3

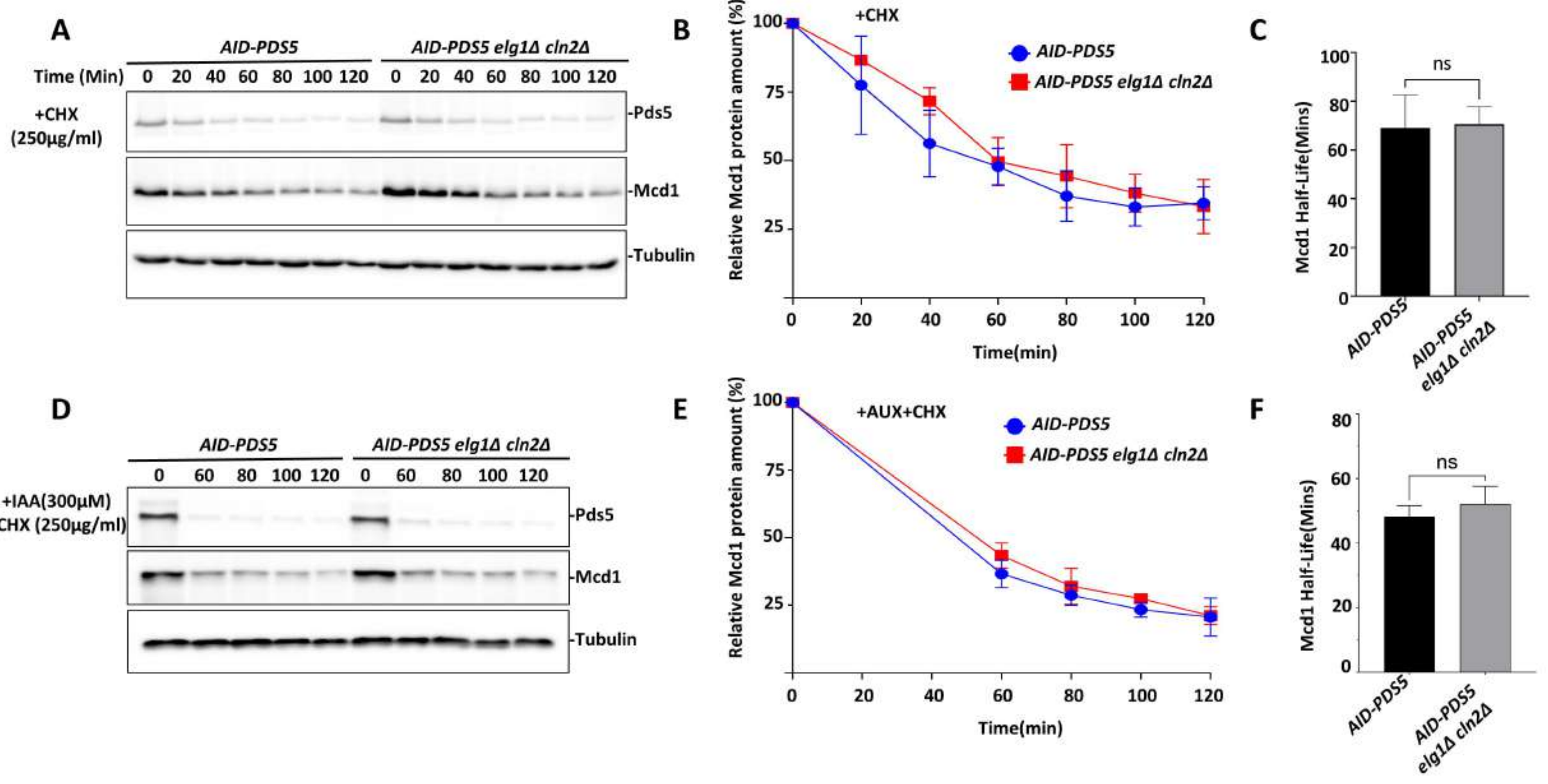
